# Molecular Determinants of Substrate Selectivity of a Pneumococcal Rgg-regulated Peptidase-Containing ABC Transporter

**DOI:** 10.1101/783472

**Authors:** Charles Y Wang, Jennifer S. Medlin, Don R. Nguyen, W. Miguel Disbennett, Suzanne Dawid

**Affiliations:** University of Michigan Department of Microbiology and immunology; University of Michigan Department of Pediatrics

**Author notes:** Don R. Nguyen currently at Michigan State University. W. Miguel Disbennett currently at Ohio State University.

## Abstract

Peptidase-containing ABC transporters (PCATs) are a widely distributed family of transporters which secrete double-glycine (GG) peptides. In the opportunistic pathogen *Streptococcus pneumoniae* (pneumococcus), the PCATs ComAB and BlpAB have been shown to secrete quorum-sensing pheromones and bacteriocins related to the competence and pneumocin pathways. Here, we describe another pneumococcal PCAT, RtgAB, encoded by the *rtg* locus and found intact in 17% of strains. The Rgg/SHP-like quorum sensing system RtgR/S, which uses a peptide pheromone with a distinctive Trp-X-Trp motif, regulates expression of the *rtg* locus and provides a competitive fitness advantage in a mouse model of nasopharyngeal colonization. RtgAB secretes a set of co-regulated *rtg* GG peptides. ComAB and BlpAB, which share a substrate pool with each other, do not secrete the *rtg* GG peptides. Similarly, RtgAB does not efficiently secrete ComAB/BlpAB substrates. We examined the molecular determinants of substrate selectivity between ComAB, BlpAB, and RtgAB and found that the GG peptide signal sequences contain all the information necessary to direct secretion through specific transporters. Secretion through ComAB and BlpAB depends largely on the identity of four conserved hydrophobic signal sequence residues previously implicated in substrate recognition by PCATs. In contrast, a motif situated at the N-terminal end of the signal sequence, found only in *rtg* GG peptides, directs secretion through RtgAB. These findings illustrate the complexity in predicting substrate-PCAT pairings by demonstrating specificity that is not dictated solely by signal sequence residues previously implicated in substrate recognition.

**Importance:** The export of peptides from the cell is a fundamental process carried out by all bacteria. One method of bacterial peptide export relies on a family of transporters called peptidase-containing ABC transporters (PCATs). PCATs export so-called GG peptides which carry out diverse functions, including cell-to-cell communication and inter-bacterial competition. In this work, we describe a PCAT-encoding genetic locus, *rtg*, in the pathogen *Streptococcus pneumoniae* (pneumococcus). The *rtg* locus is linked to increased competitive fitness advantage in a mouse model of nasopharyngeal colonization. We also describe how the *rtg* PCAT preferentially secretes a set of co-regulated GG peptides but not GG peptides secreted by other pneumococcal PCATs. These findings illuminate a relatively understudied part of PCAT biology: how these transporters discriminate between different subsets of GG peptides. Ultimately, expanding our knowledge of PCATs will advance our understanding of the many microbial processes dependent on these transporters.

## Introduction

Export of polypeptides from their site of synthesis in the cytoplasm to the extracellular space is a fundamental physiological function of all cells. The secretome, the collection of all non-membrane associated proteins secreted from the cell, may comprise up to 20% of an organism’s total proteome (1). Bacteria have evolved many different strategies for exporting proteins and peptides (2). One such strategy is the secretion of peptides using a family of ATP-binding cassette (ABC) transporters called peptidase-containing ABC transporters (PCATs).

PCATs are ABC transporters that contain characteristic N-terminal peptidase domains (PEP). PEP belongs to the family of C39 cysteine proteases and is responsible for the proteolytic processing of substrates during transport (3). In gram-positive bacteria, PCATs function either alone or with a single additional accessory protein (4). The most common function of PCATs is to assist in the biosynthesis of bacteriocins: antimicrobial peptides produced by bacteria to kill or otherwise inhibit the proliferation of other, usually closely related, bacteria (5). Some PCATs also promote cell-to-cell communication by secreting the peptide pheromones of gram-positive quorum-sensing systems (6–10). In short, PCATs are widely distributed peptide transporters which play key roles in shaping how bacteria interact with each other.

Oftentimes, expression of PCATs is under the control of quorum-sensing systems. These regulatory systems rely on cell-to-cell signaling to induce and coordinate the expression of their target genes under specific conditions. One such mode of PCAT regulation is the Rgg/SHP pathway (11). Rgg is a family of transcription regulators found in many gram-positive bacteria. In the genus Streptococcus, Rgg-family regulators are sometimes associated with short, hydrophobic peptides (SHPs) (12). SHPs are small peptides which are exported by the PptAB transporter (13, 14) and processed into mature pheromones. The Ami oligopeptide importer then internalizes the pheromones back into the cell, where they bind to and modulate the activity of Rgg-family regulators (12). Besides PCATs and bacteriocin production, Rgg/SHP systems have been found to regulate diverse processes such as carbohydrate utilization (15), tissue invasion (13, 16), capsule production (16, 17), and biofilm formation (17, 18). A related group of Rgg regulators, the ComRs, are associated with SHP-like pheromones called ComS or XIP (SigX-inducing peptide) and control competence activation in some streptococcal species (19, 20).

In the gram-positive opportunistic pathogen *Streptococcus pneumoniae* (pneumococcus), the PCATs ComAB and BlpAB secrete quorum-sensing pheromones that control two important cellular pathways: genetic competence (the ability to take up and incorporate extracellular DNA into the genome) and production of the major family of pneumococcal bacteriocins (Blp bacteriocins, or pneumocins) (6, 7, 21, 22). ComAB and BlpAB secrete the same GG peptides, including the competence- and pneumocin-inducing pheromones and the pneumocins (23–25). Substrate sharing between ComAB and BlpAB affects competence and pneumocin regulation and influences when and with what effectiveness naturally occurring BlpAB^+^ and BlpAB^−^ strains can employ pneumocin-mediated killing (25, 26).

The functional implications of the shared ComAB/BlpAB substrate pool highlight the need to better understand how different PCATs select their substrates. PCAT substrates contain N-terminal signal sequences (also called leader peptides) which terminate in a Gly-Gly (sometimes also Gly-Ala or Gly-Ser) motif (3). For this reason, they are referred to as double-glycine (GG) peptides. During transport, PEP cleaves the peptide bond following the GG motif to remove the signal sequence from the C-terminal mature peptide fragment (cargo peptide). The signal sequences of GG peptides bind to PEP of PCATs through hydrophobic interactions involving three or four conserved residues in the signal sequences (27–29). These residues are located at positions −4, −7, −12, and −15 relative to the scissile bond. The GG motif allows the substrate to fit in the narrow entrance to the active site of PEP and is also required for binding and cleavage (28, 30, 31). Besides these conserved residues, the signal sequences of different GG peptides are fairly divergent. Mutagenesis studies of several different PCAT-substrate pairs have largely failed to identify any contribution of these non-conserved residues to substrate-PEP binding (27–29).

While substantial progress has been made in uncovering the mechanisms that allow PCATs to recognize GG peptides, comparatively little is known about how or if PCATs discriminate between different GG peptides. In addition to ComAB and BlpAB from pneumococcus, multiple PCATs have been shown to process and/or secrete multiple peptides with distinct signal sequences, sometimes even those from different strains or species (28, 32–35). These data suggest that in general, PCATs are not particularly selective when it comes to choosing substrates.

In this work, we describe a previously uncharacterized locus in pneumococcus, *rtg*, which encodes the PCAT RtgAB and several GG peptides. This locus is regulated by the Rgg/SHP-like system RtgR/S, which provides a competitive fitness advantage during nasopharyngeal colonization. We demonstrate that RtgAB secretes the *rtg* GG peptides but not ComAB/BlpAB substrates, nor can ComAB or BlpAB efficiently secrete the *rtg* GG peptides. Finally, we investigate the signal sequence determinants that selectively direct peptides toward either RtgAB or ComAB/BlpAB and show that a unique N-terminal motif is required for secretion by RtgAB. These findings shed light on how PCATs can use signal sequence motifs beyond the previously described conserved hydrophobic residues to distinguish different GG peptides.

## Results

### Identification of an uncharacterized pneumococcal PCAT-encoding locus

As part of an effort to catalog the PCAT repertoire of *S. pneumoniae*, we searched pneumococcal genomes for putative PCAT genes that had not been previously described. One of the hits was *CGSSp9BS68_07257* (henceforth *07257*), a gene found in the clinical isolate Sp9-BS68 (36) (Fig. 1A). Upstream of *07257* is a gene oriented in the opposite direction predicted to encode an Rgg-family transcription regulator (12). We hypothesized that this regulator controls expression of *07257* and named the locus *rtg* (<u>R</u>gg-regulated transporter of double-<u>g</u>lycine peptides), the transporter gene *rtgA* and the regulator gene *rtgR*. *rtgR* marks one end of the locus and is separated from a partially disrupted arginine biosynthesis cluster (37) by two transcription terminators. Downstream of *rtgA* are several genes arranged in a single operon. These include *rtgB*, which encodes a putative ComB/BlpB-like transport accessory protein, and the GG peptide genes *rtgG*, *rtgT*, *rtgW1*, and *rtgW2*. A transcription terminator separates the last gene, *rtgD2*, from a disrupted putative endoRNase gene and *pspA*. A different version of the *rtg* locus is found in the laboratory strain D39 (and its derivative, R6) but with a disrupted *rtgA* (Fig. 1A).

**Figure 1.**
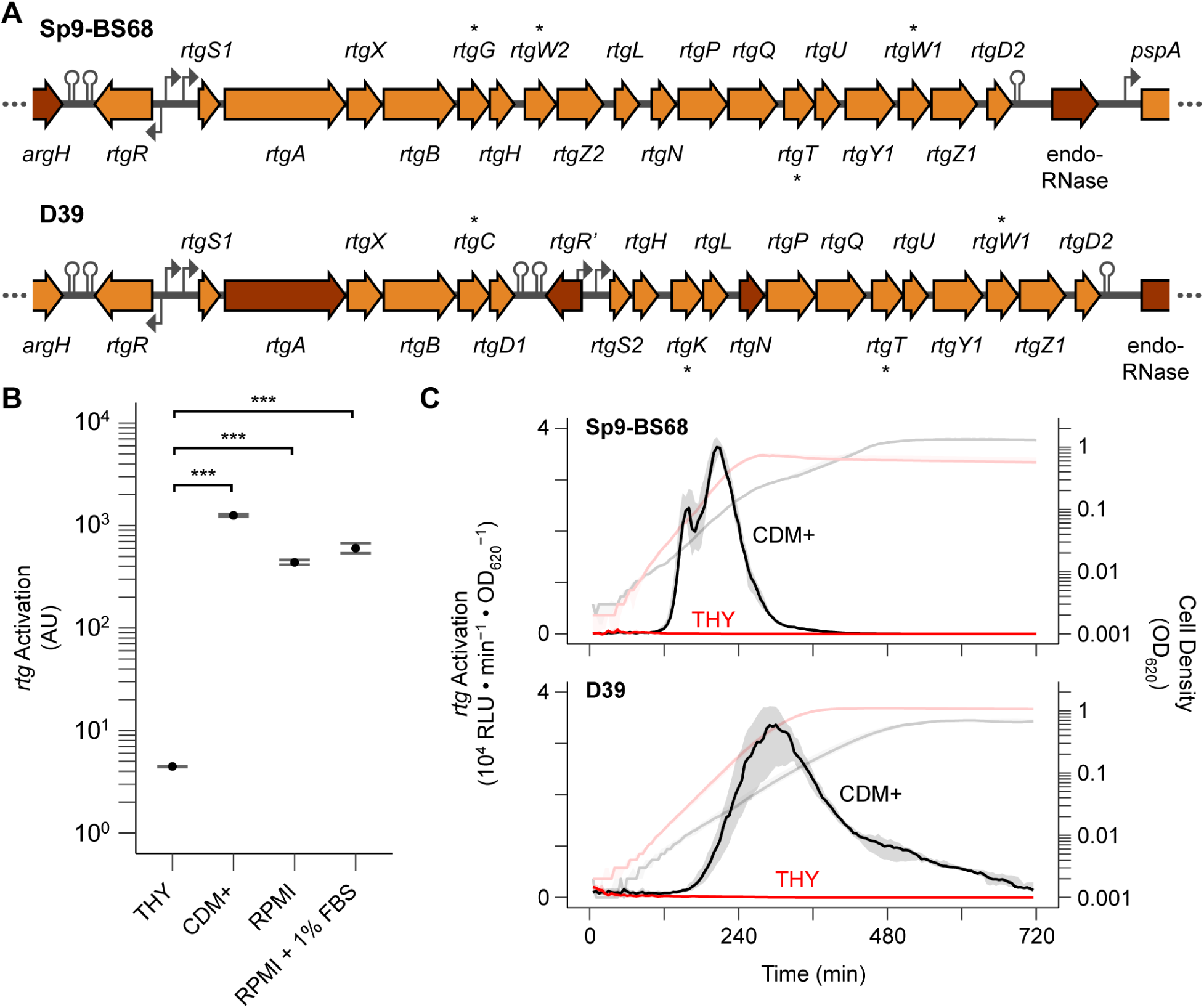
*rtg* is an actively regulated PCAT-encoding locus in pneumococcus. **(A)** Organization of *rtg* in strains Sp9-BS68 and D39. Block arrows represent genes. Dark shading indicates a pseudogene. Bent arrows represent promoters. Hairpins represent transcription terminators. GG peptide genes are marked with asterisks. **(B)** Activation of *rtg* in various growth media. A Sp9-BS68 P*_rtgA_*-*luc* reporter strain was grown in the indicated media, and luciferase activity was sampled when cells reached OD_620_ 0.2. The plotted values were normalized against cell density and the luciferase activity from a strain harboring constitutively expressed luciferase grown in the same medium. Plots show mean ± S.E. of 3 independent experiments. Statistics: *** p < 0.001; ANOVA with Tukey’s HSD. **(C)** Timing of *rtg* activation in THY and CDM+. Sp9-BS68 (top) and D39 (bottom) P*_rtgS1_*-*luc* reporter strains were grown in THY (red) and CDM+ (black) and monitored for *rtg* activation (dark, left y axis) and cell density (light, right y axis). Plots show median (line) and 25% to 75% quantiles (shading) of 12 wells pooled from 3 independent experiments.

### The Rgg/SHP-like pheromone pair RtgR/RtgS regulates rtg

We found that *rtg* expression is inhibited in mid-exponential phase during growth in the peptide-rich medium THY (Todd-Hewitt broth + 0.5% yeast extract) but highly upregulated in the peptide-poor media CDM+ (38) and RPMI (Fig. 1B). The start of *rtg* activation in both Sp9-BS8 and D39 during growth in CDM+ occurs in early exponential phase at cell densities as low as OD_620_ 0.01 (Fig. 1C). In contrast, *rtg* stays inactive in THY throughout the exponential and stationary phases. We concluded from these data that *rtg* is actively regulated, most likely by RtgR. Since Rgg regulators are often associated with peptide pheromones, we searched for and found an ORF in Sp9-BS68 between *rtgR* and *rtgA* predicted to encode a SHP-like pheromone. D39 has two copies of the candidate pheromone: one located between *rtgR* and *rtgA* and the other downstream of *rtgB*. We named the only copy of the ORF in Sp9-BS68 and the first copy in D39 (between *rtgR* and *rtgA*) *rtgS1* and the second copy in D39 *rtgS2* (Fig. 1A).

Having identified a putative Rgg/SHP-like regulatory system, we sought to define the contributions of RtgR and RtgS to *rtg* regulation through deletional analysis. We monitored *rtg* activation in Sp9-BS68 Δ*rtgS1*, Δ*rtgR*, and Δ*rtgS1*Δ*rtgR* strains during growth in CDM+ and THY (Fig. 2A). None of the mutants showed signs of *rtg* activation in either medium, indicating that both RtgR and RtgS promote and are required for *rtg* activation. In D39, *rtgS1* encodes a peptide with a single amino acid change (S14L) compared to RtgS from Sp9-BS68, and *rtgS2* encodes a peptide with a different single amino acid change (P27S) (Fig. 2B). We found that the D39 Δ*rtgS1* and Δ*rtgS1*Δ*rtgS2* mutants failed to activate *rtg* in CDM+ while the Δ*rtgS2* mutant was indistinguishable from the wild-type strain (Fig. 2C). This indicated that the P27S substitution in the *rtgS2* product prevents it from activating *rtg*, while the S14L substitution in the *rtgS1* product does not appreciably affect signaling. Therefore, we classified the *rtgS1* product in both Sp9-BS68 and D39 as type A pheromone (RtgS_A_) and the *rtgS2* product in D39 as type B (RtgS_B_).

**Figure 2.**
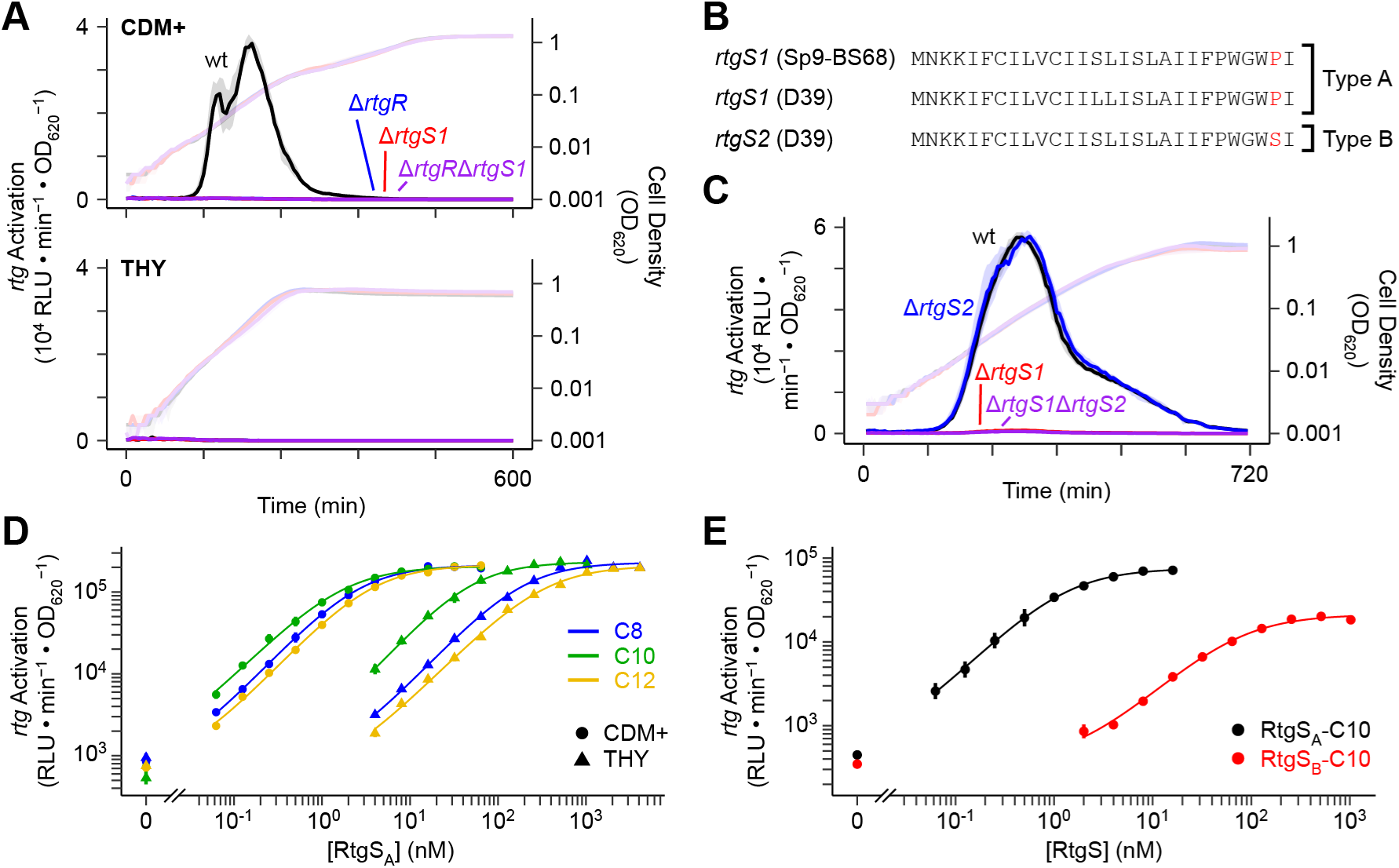
The Rgg/SHP-like RtgR/RtgS system regulates *rtg*. **(A)** *rtgR* and *rtgS1* are required for *rtg* activation in Sp9-BS68. Sp9-BS68 P*_rtgS1_*-*luc* reporters were grown in CDM+ or THY and monitored for *rtg* activation (dark, left y axis) and cell density (light, right y axis). Plots show median (line) and 25% to 75% quantiles (shading) of 12 wells pooled from 3 independent experiments. **(B)** Translated *rtgS* gene products from Sp9-BS68 and D39. The type-defining residue is highlighted in red. **(C)** *rtgS1* but not *rtgS2* is required for *rtg* activation in D39. D39 P*_rtgS1_*-*luc* reporters were grown in CDM+ and monitored for *rtg* activation (dark, left y axis) and cell density (light, right y axis). Plots show median (line) and 25% to 75% quantiles (shading) of 30 wells pooled from 3 independent experiments. **(D)** C-terminal fragments of RtgS_A_ induce *rtg*. A Sp9-BS68 Δ*rtgS1* P*_rtgS1_*-*luc* reporter was grown in CDM+ or THY to OD_620_ 0.02 and treated with synthetic RtgS_A_ fragments. Response was defined as the maximum observed P*_rtgS1_* activity within 60 min of treatment. Plotted data points represent mean ± S.E. of 3 independent experiments. **(E)** RtgS_B_ is a partial agonist at the *rtg* locus. A D39 Δ*rtgS1*Δ*rtgS2* P*_rtgS1_*-*luc* reporter was grown in CDM+ to OD_620_ 0.02 and treated with synthetic RtgS fragments. Response was defined as the maximum observed P*_rtgS1_* activity within 60 min of treatment. Plotted data points represent mean ± S.E. of 3 independent experiments.

To confirm that RtgS is the specific pheromone inducer of *rtg*, we performed dose-response assays using synthetic peptides corresponding to the C-terminal eight, ten, and twelve residues of RtgS_A_ (RtgS_A_-C8, RtgS_A_-C10, RtgS_A_-C12, respectively). All three synthetic peptides induced expression from the Sp9-BS68 *rtgS1* promoter in both CDM+ and THY, though the curves for the latter were shifted to the right in a manner consistent with pure competitive inhibition (Fig. 2D, Table 1). We also confirmed that *rtg* induction by synthetic RtgS requires RtgR (Fig. S1). In D39, RtgS_A_-C10 induces *rtg* similarly to Sp9-BS68 whereas RtgS_B_-C10 acts as a partial agonist with a 55-fold larger EC50 value than RtgS_A_-C10 (Fig. 2E, Table 1). Therefore, the Pro to Ser substitution in RtgS_B_ interferes with signaling at a step following pheromone secretion. Although partial agonists can act as competitive antagonists of full agonists, we did not observe an inhibitory phenotype associated with RtgS_B_ during natural *rtg* activation (Fig. 2C) and competitive dose-response assays showed that RtgS_B_-C10 only antagonizes RtgS_A_-C10 at likely supraphysiological concentrations (≥ 256 nM) (Fig. S2).

**Table 1.** RtgS dose-response parameters.

| Strain | Growth Medium | Peptide | EC50 (nM)<sup>a</sup> | Maximal response (10<sup>3</sup> RLU · min<sup>-1</sup> · OD<sub>620</sub><sup>-1</sup>)<sup>a</sup> |
| --- | --- | --- | --- | --- |
| Sp9-BS68<sup>b</sup> | CDM+ | RtgS<sub>A</sub>-C8 | 2.53 ± 0.25 | 212.2 ± 9.4 |
|  |  | RtgS<sub>A</sub>-C10 | 1.63 ± 0.19 | 205.7 ± 9.1 |
|  |  | RtgS<sub>A</sub>-C12 | 3.51 ± 0.28 | 213.9 ± 8.0 |
|  | THY | RtgS<sub>A</sub>-C8 | 184 ± 15 | 229.7 ± 8.6 |
|  |  | RtgS<sub>A</sub>-C10 | 47.4 ± 6.9 | 232.4 ± 14.3 |
|  |  | RtgS<sub>A</sub>-C12 | 315 ± 42 | 210.1 ± 13.1 |
| D39<sup>c</sup> | CDM+ | RtgS<sub>A</sub>-C10 | 1.25 ± 0.30 | 75.1 ± 8.1 |
|  |  | RtgS<sub>B</sub>-C10 | 68.8 ± 11.5 | 21.2 ± 1.5 |

<sup>a</sup> Mean ± S.E.; derived from fitting data to Hill model.<sup>b</sup> $\Delta$ rtgS1<sup>c</sup> $\Delta$ rtgS1 $\Delta$ rtgS2**Table 2.** Genes of the *rtg* locus.

Next, we determined that, consistent with previously described Rgg/SHP systems, *rtg* activation requires both the Ami importer and the PptAB transporter (Fig. S3A). We also showed that response to exogenous RtgS treatment requires Ami but not PptAB (Fig. S3B), consistent with their respective roles as pheromone importer and exporter.

Finally, mutagenesis of the Sp9-BS68 *rtgS1* promoter region revealed the presence of two nearly identical promoters, each contributing partially to *rtgS1* expression (Fig. S4A,B). We also identified an inverted repeat found in both promoters which is required for RtgS-induced expression and likely represents the RtgR binding site (Fig. S4A,C).

### RtgAB secretes rtg-encoded GG peptides

Sp9-BS68 and D39 both encode four putative GG peptides at the *rtg* locus (Fig. 1A). To determine whether RtgAB secretes the *rtg* GG peptides, we employed a HiBiT tag-based peptide secretion assay (25). We constructed autoinducing-deficient Δ*rtgS1*Δ*rtgS2* R6 strains which encode HiBiT-tagged *rtgC* (from D39) or *rtgG* (from Sp9-BS68) placed downstream of *rtgB* in RtgAB^−^ (native R6 *rtgA* pseudogene) and RtgAB^+^ (pseudogene-repaired) backgrounds. These strains were also ComAB^−^ and BlpAB^−^ in order to remove the possibility of peptide secretion through these other PCATs. Upon RtgS-C10 treatment in CDM+, levels of extracellular RtgC-HiBiT and RtgG-HiBiT in the RtgAB^+^ cultures increase to 26- and 376-fold, respectively, compared to their levels in the RtgAB^−^ cultures (Fig. 3). From these data, we conclude that RtgAB secretes the *rtg* GG peptides.

**Figure 3.**
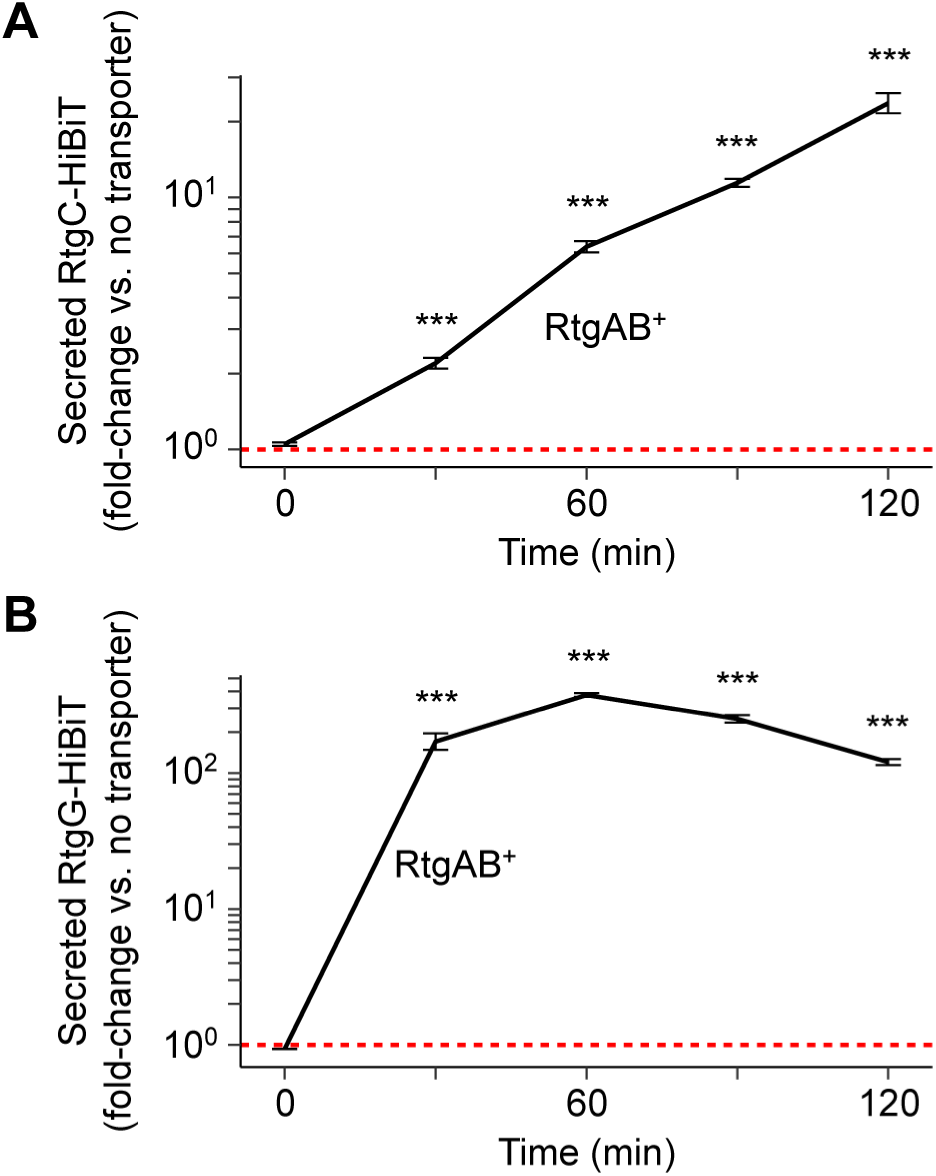
RtgAB secretes RtgC and RtgG. **(A-B)** R6 ComAB^−^/BlpAB^−^/RtgAB^−^ and ComAB^−^/BlpAB^−^/RtgAB^+^ strains expressing RtgC-HiBiT (A) or RtgG-HiBiT (B) were grown in CDM+ to OD_620_ 0.05 and treated with 200 ng/mL CSP, 200 ng/mL BlpC, and 20 nM RtgS_A_-C10. Samples were taken every 30 min and extracellular HiBiT signal was quantified. Data are presented as fold-change values relative to the ComAB^−^/BlpAB^−^/RtgAB^−^ control. Red, dashed line represents fold-change = 1 (no difference vs. the control). Plots show mean ± S.E. of 3 independent experiments. Statistics: comparisons vs. ComAB^−^/BlpAB^−^/RtgAB^−^ at each timepoint; *** p < 0.001; ANOVA with Tukey’s HSD.

### The rtg locus exhibits extensive variation across different pneumococcal strains

In order to catalog the diversity found at *rtg*, we conducted a survey of the locus in a collection of pneumococcal clinical isolates from Massachusetts, USA (39). After removing genomes in which *rtg* spans multiple contigs or when conserved genes flanking *rtg* could not be found, we were left with 318 out of 616 strains, all of which encoded at least one *rtg* gene. We analyzed the *rtg* loci from these 318 strains and clustered them into 23 groups based on overall architecture (Fig. S5). Across all 318 loci, we found 24 unique *rtg* genes (Table 2). Searching for these genes in the full collection revealed that 615 out of 616 strains had some version of the *rtg* locus. Next, we analyzed the variation in RtgS among the 318 filtered strains. We found at least one copy of *rtgS* in each strain. Because duplication of *rtgS* is common, we assigned the name *rtgS1* to any copy located next to *rtgR* and the name *rtgS2* to any copy located next to *rtgR’*. Based on the identity of the penultimate residue in the translated peptide, which we have shown is important for signaling activity, we catalogued a total of three pheromone types: types A (Pro), B (Ser), and C (Leu). Only two other positions in the last 12 residues of RtgS show variation: Ala/Val at position −10 from the C-terminus and Ile/Val at position −8. The functional significance of these other polymorphisms is unknown. Finally, we analyzed each strain’s RtgAB status and found 5% of strains carry unambiguously intact *rtgA* and *rtgB*. Another 12% carry intact *rtgB* and a version of *rtgA* with a start codon mutation (ATG>ATT) but is otherwise intact. We determined that a strain with the ATG>ATT mutation still produces functional RtgAB, likely by using an alternative start site, and suffers only a minor reduction in secretion capacity compared to a strain with fully intact *rtgA* (Fig. S6). Therefore, 17% of strains encode a functional RtgAB.

**Table 2.** Genes of the *rtg* locus.

| Gene |  | Product | Frequency<sup>a,b</sup> |
| --- | --- | --- | --- |
| *rtgA* | Intact | Peptidase-containing ABC transporter | 5.0% |
|  | Start codon mutation only | Peptidase-containing ABC transporter | 14.5% |
|  | Disrupted | Peptidase-containing ABC transporter, truncated | 59.1% |
| *rtgB* | Intact | Putative transport accessory protein | 18.6% |
|  | Disrupted | Putative transport accessory protein, truncated | 2.5% |
| *rtgC* |  | GG peptide | 15.1% |
| *rtgD* |  | Hypothetical protein | 99.7% |
| *rtgE* |  | GG peptide | 7.2% |
| *rtgG* |  | GG peptide | 4.1% |
| *rtgH* |  | Hypothetical protein | 40.6% |
| *rtgK* |  | GG peptide | 41.2% |
| *rtgL* |  | Hypothetical protein | 83.3% |
| *rtgM* |  | GG peptide | 59.7% |
| *rtgN* |  | Hypothetical protein | 82.4% |
| *rtgP* |  | Putative integral membrane protein | 83.3% |
| *rtgQ* |  | Putative integral membrane protein | 83.3% |
| *rtgR* |  | Rgg-family transcription regulator | 78.6% |
| *rtgR'* |  | Rgg-family transcription regulator, truncated | 36.5% |
| *rtgS* | Type A | SHP-like pheromone, type A | 42.5% |
|  | Type B | SHP-like pheromone, type B | 67.6% |
|  | Type C | SHP-like pheromone, type C | 0.6% |
| *rtgT* |  | GG peptide | 83.0% |
| *rtgU* |  | Hypothetical protein | 83.0% |
| *rtgV* |  | GG peptide; probable fusion of RtgW and RtgZ | 35.5% |
| *rtgW* |  | GG peptide | 83.0% |
| *rtgX* |  | GyrI-like hypothetical protein | 21.1% |
| *rtgY* |  | Putative integral membrane protein | 88.7% |
| *rtgZ* |  | Putative integral membrane protein | 99.7% |

<sup>a</sup> In 318 genomes from the Massachusetts collection with fully sequenced, gap-less *rtg* loci.<sup>b</sup> May include pseudogenes unless stated otherwise.

### Active RtgR/S confers a competitive fitness advantage during nasopharyngeal colonization

In order to determine the biological role of *rtg*, we tested the effect of a regulatory deletion on colonization of the nasopharynx, the natural niche of pneumococcus. Despite similar levels of colonization between the wildtype and Δ*rtgR*Δ*rtgS1* strains in singly inoculated mice at 3 days post-inoculation (Fig. 4A), the wildtype strain outcompeted the mutant in co-inoculated mice (Fig. 4B). These data suggest that RtgR/S is active during nasopharyngeal colonization and show that active RtgR/S provides a fitness advantage over RtgR/S-inactive strains during co-colonization.

**Figure 4.**
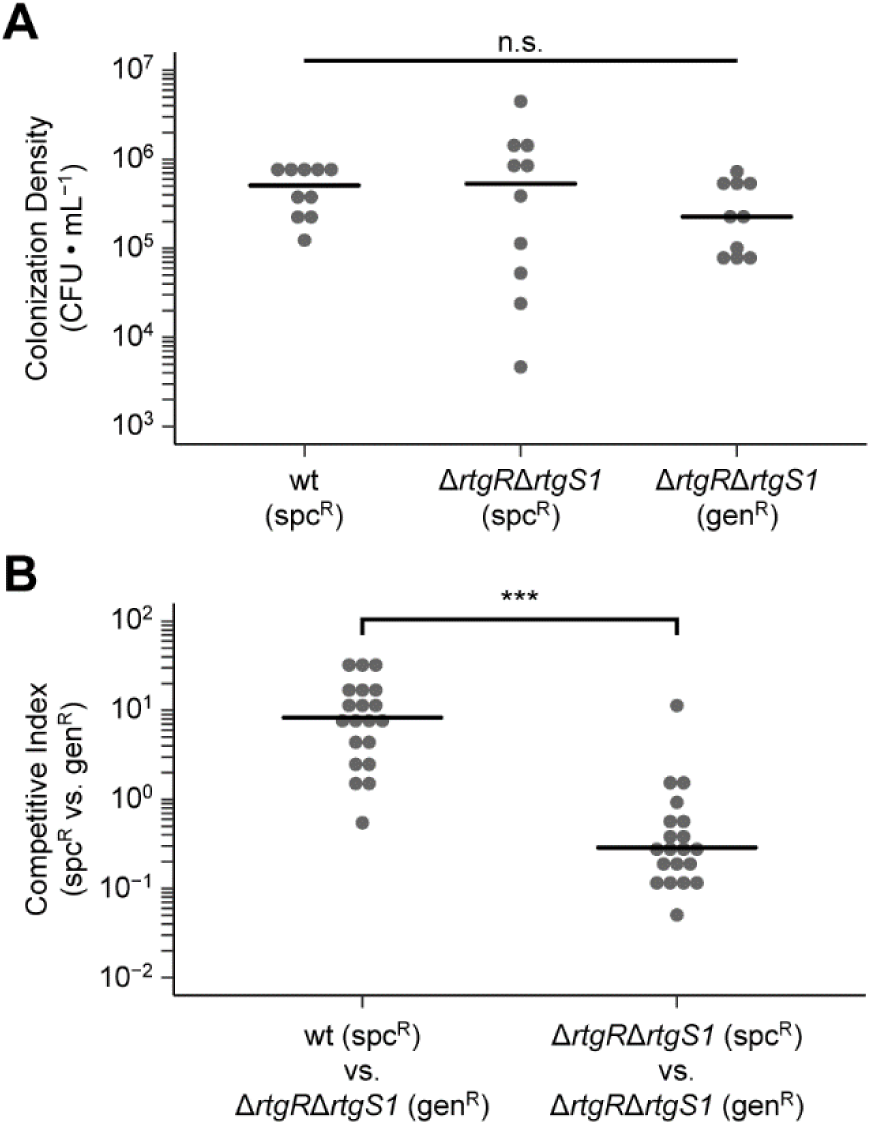
Active RtgR/S provides a competitive fitness advantage during nasopharyngeal colonization. **(A)** Sp9-BS68 strains were singly inoculated into the nasopharynx of 5- to 7-week-old female BALB/c mice. At 3 days post-inoculation, colonization density was sampled by nasal wash. Black bars represent medians. N = 10 mice per strain pooled from 2 independent experiments. Statistics: n.s. not significant; Kruskal-Wallis test. **(B)** Pairs of Sp9-BS68 strains were co-inoculated into the nasopharynx of 5- to 7-week-old female BALB/c mice. At 3 days post-inoculation, colonization density was sampled by nasal wash and competitive indices were calculated. Black bars represent medians. N = 20 mice per competition pooled from 2 independent experiments. Statistics: *** p < 0.001; Mann-Whitney test.

### RtgAB and ComAB/BlpAB preferentially secrete different sets of peptides

The pneumococcal PCATs ComAB and BlpAB secrete the same diverse set of GG peptides (23–25). Therefore, we wondered if ComAB and BlpAB could also secrete the *rtg* GG peptides and if RtgAB is similarly promiscuous and could secrete ComAB/BlpAB substrates. We repeated the RtgC-HiBiT and RtgG-HiBiT secretion assays with ComAB^+^ and BlpAB^+^ strains, using treatment with the *com* and *blp* pheromones CSP and BlpC, respectively, to induce their expression. ComAB and BlpAB secrete markedly reduced amounts of RtgC-HiBiT and RtgG-HiBiT compared to RtgAB (Figs. 5A,B). To determine if RtgAB could secrete a ComAB/BlpAB substrate, we assayed secretion of a HiBiT tagged version of the BlpI bacteriocin driven by its native promoter. RtgAB secretes roughly ten-fold less BlpI-HiBiT than BlpAB (Fig. 5C). Therefore, RtgAB and ComAB/BlpAB do not efficiently cross-secrete each other’s substrates. Consistent with this, RtgAB^+^ strains do not show differences in *com* or *blp* activation compared to RtgAB^−^ strains during growth in CDM+ (Fig. S7). Under these conditions *rtg* is expected to turn on before *com* or *blp* in every strain background. Thus, even when RtgAB is highly expressed it secretes too little CSP and BlpC to affect the timing of *com* and *blp* activation.

**Figure 5.**
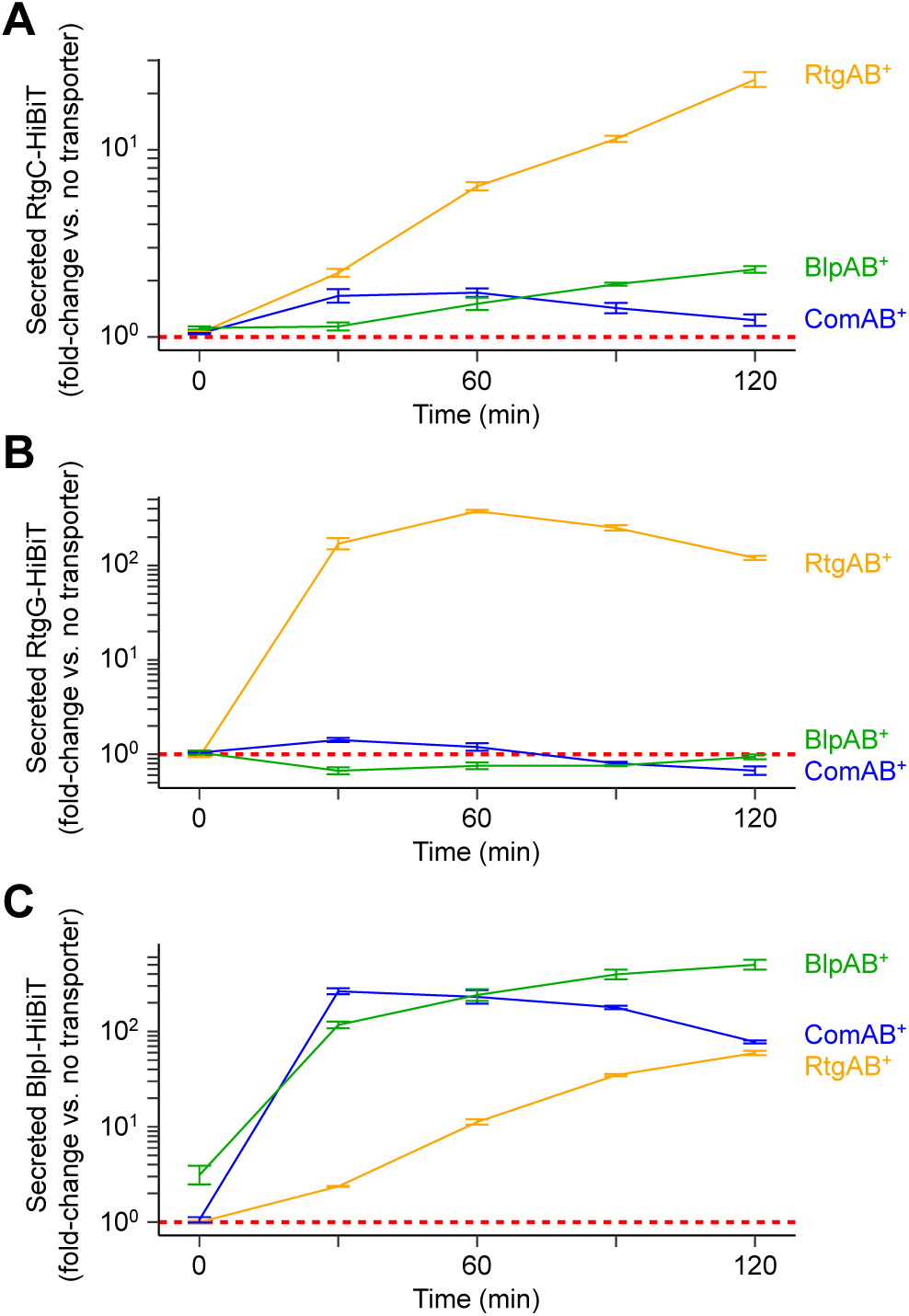
RtgAB secretes different GG peptides from ComAB and BlpAB. **(A-C)** R6 ComAB^−^/BlpAB^−^/RtgAB^−^ and single-transporter positive ComAB^+^, BlpAB^+^, RtgAB^+^ strains expressing RtgC-HiBiT (A), RtgG-HiBiT (B), or BlpI-HiBiT (C) were grown in CDM+ to OD_620_ 0.05 and treated with 200 ng/mL CSP, 200 ng/mL BlpC, and 20 nM RtgS_A_-C10. Samples were taken every 30 min and extracellular HiBiT signal was quantified. Data are presented as fold-change values relative to the ComAB^−^/BlpAB^−^/RtgAB^−^ control. Red, dashed line represents fold-change = 1 (no difference vs. the control). Plots show mean ± S.E. of 3 independent experiments.

### RtgAB and ComAB/BlpAB recognize their substrates through different signal sequence motifs

Given that we had found RtgAB and ComAB/BlpAB do not share the same substrate pool, we explored how the transporters discriminate between substrate and non-substrate GG peptides. We showed that the BlpI signal sequence (SS_BlpI_) prevents secretion of the RtgG cargo peptide through RtgAB (Fig. 6A). However, it did not promote secretion of RtgG through ComAB/BlpAB, suggesting an incompatibility between the cargo peptide and these two transporters. On the other hand, the RtgG signal sequence (SS_RtgG_) both promotes secretion of the BlpI cargo peptide through RtgAB while preventing its secretion through ComAB and BlpAB (Fig. 6B).

**Figure 6.**
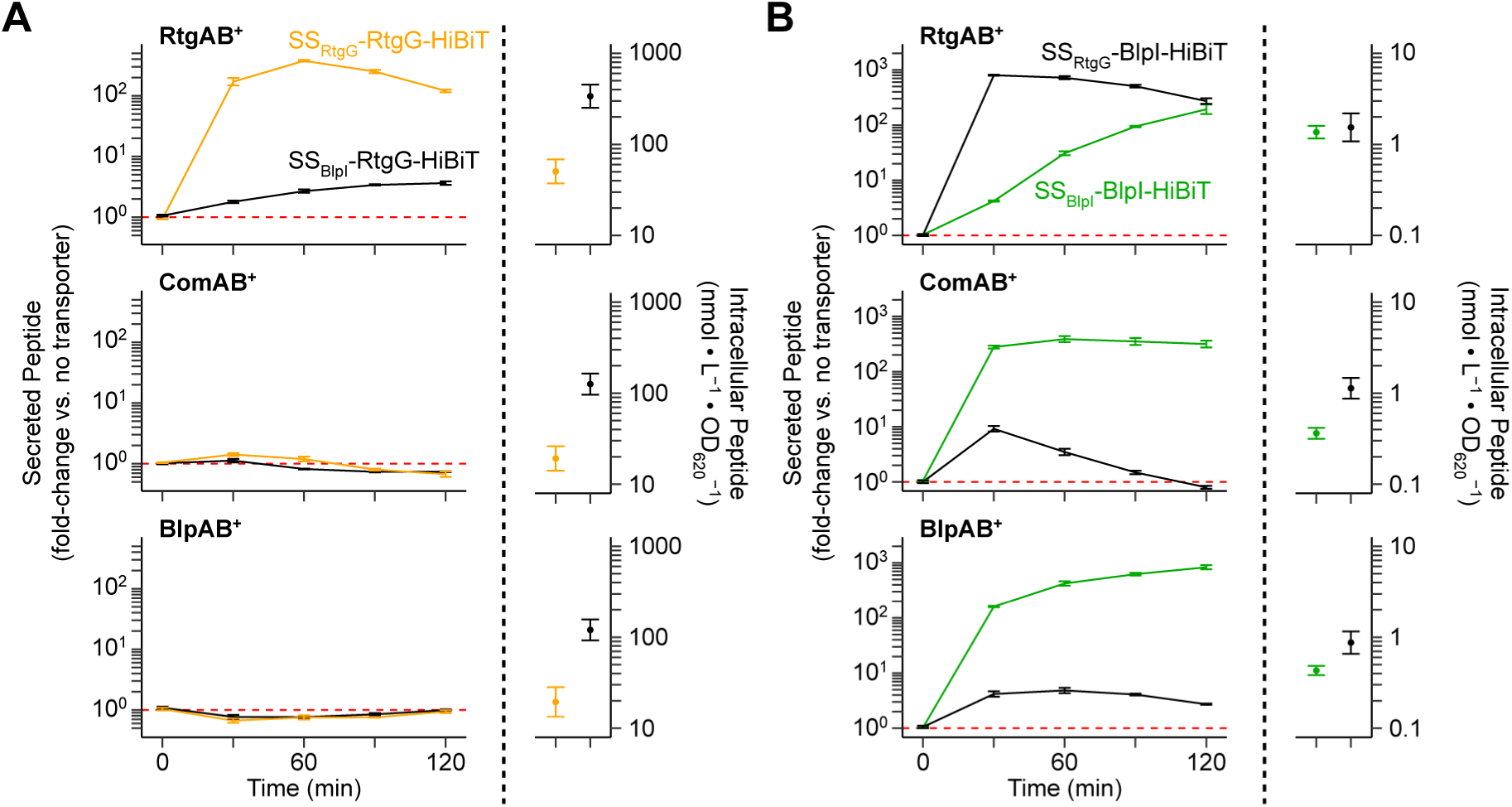
RtgAB, ComAB, and BlpAB recognize GG peptides through their N-terminal signal sequences. **(A-B)** R6 ComAB^−^/BlpAB^−^/RtgAB^−^ and single-transporter positive ComAB^+^, BlpAB^+^, RtgAB^+^ strains expressing signal sequence-swapped RtgG-HiBiT cargo peptide (A), or BlpI-HiBiT cargo peptide (B) were grown in CDM+ to OD_620_ 0.05 and treated with 200 ng/mL CSP, 200 ng/mL BlpC, and 20 nM RtgS_A_-C10. Samples were taken every 30 min and extracellular HiBiT signal was quantified (left). Data are presented as fold-change values relative to the ComAB^−^/BlpAB^−^/RtgAB^−^ control. Red, dashed line represents fold-change = 1 (no difference vs. the control). At the 120-min timepoint, intracellular peptide content was also quantified (right). Plots show mean ± S.E. of 3 independent experiments.

To rule out the possibility of differences in peptide expression being solely responsible for the secretion differences, we also measured the amount of intracellular peptide in each assay (Fig. 6A,B; right-hand graphs). The signal sequence swaps did affect intracellular peptide levels. However, these intracellular differences cannot account for the observed changes in secretion; higher intracellular levels did not correlate with more secretion, and while intracellular levels of the same peptide were relatively consistent across different strains (RtgAB^+^ vs. ComAB^+^ vs. BlpAB^+^), secretion was not. Thus, the observed changes in secretion between the different peptides most likely reflect differences in peptide-transporter interactions.

In conclusion, while cargo peptide can dictate transporter compatibility in some cases, the signal sequences of GG peptides still contain all the necessary information to direct secretion of their cargo peptides through the proper transporters. For all future assays, we used BlpI as the cargo peptide since it can be secreted by all three transporters given the correct signal sequence.

Next, we searched for the specific signal sequence residues involved in transport selectivity. We found that secretion of peptide through ComAB/BlpAB depends on the identities of the residues at the conserved signal sequence positions −15, −12, −7, and −4. These positions had been previously implicated in substrate recognition by PCATs (27, 28, 30). The combination of the four residues at these positions from SS_BlpI_ (F/M/L/V) introduced into SS_RtgG_ promote secretion through ComAB/BlpAB although they are not strictly required for secretion in the context of SS_BlpI_ (Fig. 7A). The complementary association does not hold for RtgAB-mediated secretion in that the four residues from SS_RtgG_ (Y/L/M/L) are neither necessary nor sufficient for secretion through RtgAB (Fig. 7A). Additionally, alanine substitutions at all four positions in SS_RtgG_ only partially impede secretion through RtgAB, but the same substitutions in SS_BlpI_ prevent secretion through ComAB and BlpAB almost entirely (Fig. 7B).

**Figure 7.**
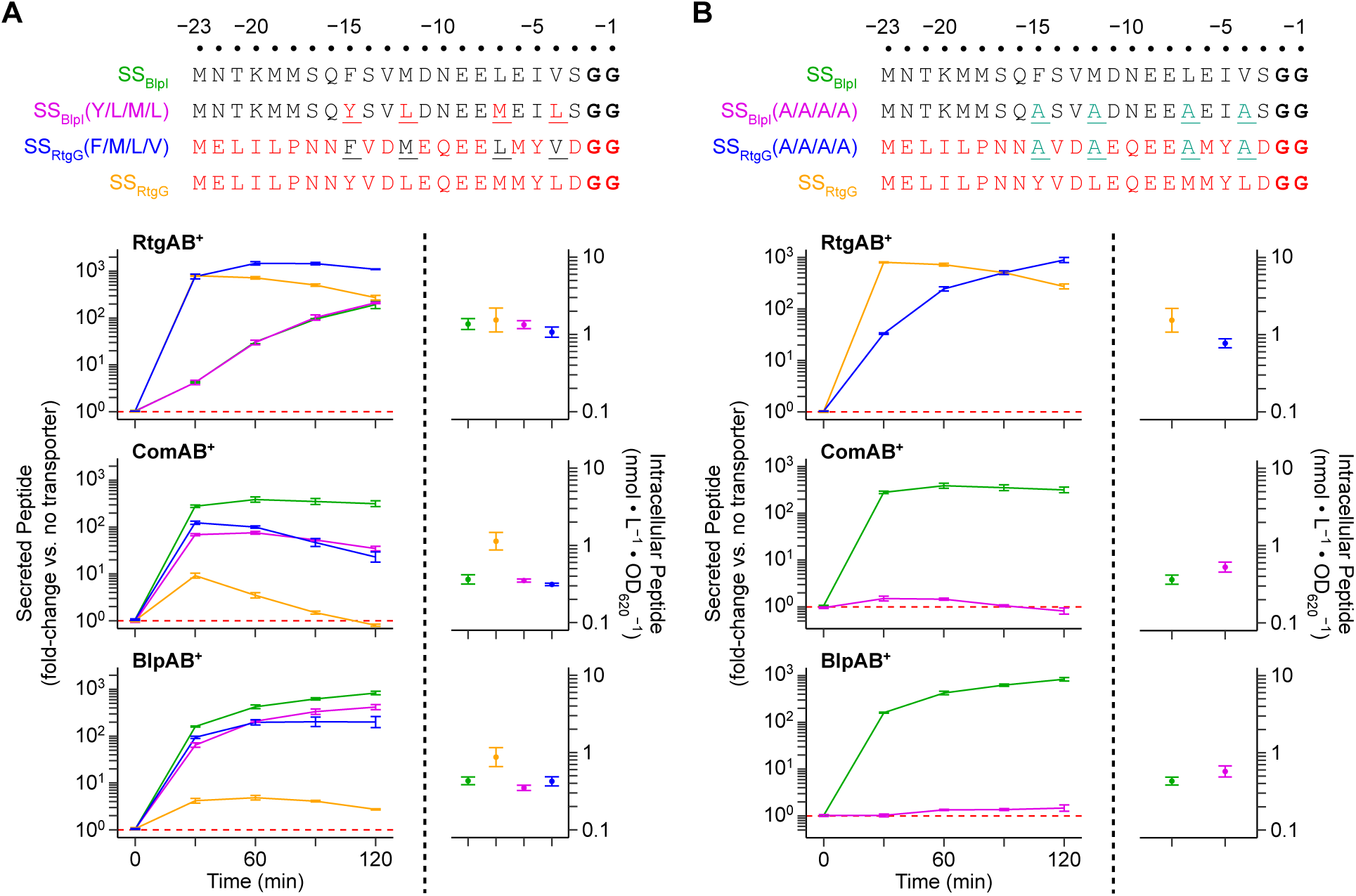
Specific GG peptide signal sequence residues at positions −15, −12, −7, and −4 modulate secretion by ComAB and BlpAB, but not RtgAB. **(A-B)** R6 ComAB^−^/BlpAB^−^/RtgAB^−^ and single-transporter positive ComAB^+^, BlpAB^+^, RtgAB^+^ strains expressing BlpI or RtgG signal sequences with residue swaps (A) or alanine substitutions (B) at positions −15, −12, −7, and −4 (top; mutated positions are underlined) fused to BlpI-HiBiT cargo peptide were grown in CDM+ to OD_620_ 0.05 and treated with 200 ng/mL CSP, 200 ng/mL BlpC, and 20 nM RtgS_A_-C10. Samples were taken every 30 min and extracellular HiBiT signal was quantified (left). Data are presented as fold-change values relative to the ComAB^−^/BlpAB^−^/RtgAB^−^ control. Red, dashed line represents fold-change = 1 (no difference vs. the control). At the 120-min timepoint, intracellular peptide content was also quantified (right). Plots show mean ± S.E. of 3 independent experiments.

### A specific motif at the N-terminal ends of rtg GG peptide signal sequences promotes secretion through RtgAB

In order to identify the signal sequence residues that promote secretion through RtgAB, we turned our attention to the N-terminal ends of the signal sequences, which are conserved in *rtg* GG peptides but not ComAB/BlpAB substrates. Residue swaps at positions −22 to −18 in SS_RtgG_ and SS_BlpI_ demonstrated that secretion through RtgAB, but not ComAB or BlpAB, depends on specific signal sequence residues in this region (Fig. 8A). The P(−18)M substitution in SS_RtgG_ modestly decreases secretion through RtgAB, and removal of all residues on the N-terminal side of this substitution further decreases secretion (Fig. 8B). Meanwhile, removal of the residues at the same positions from SS_BlpI_ did not change secretion through RtgAB (Fig. 8C). These data indicate that the residues in this region in SS_RtgG_ were selected to interact with RtgAB rather than to avoid steric clash. Alanine scanning mutagenesis of the −22 to −19 region of SS_RtgG_ revealed that secretion through RtgAB was not sensitive to mutation at any single site (Fig. S8). These data can be explained by multiple, redundant residues mediating the interactions in this region or the interactions being tolerant to alanine substitution. We conclude that RtgAB recognizes *rtg* GG peptides through interactions involving the signal sequence residues in the −22 to −18 region. At the same time, RtgAB’s substrate recognition mechanism has evolved to be less reliant than that of ComAB or BlpAB on interactions with the hydrophobic signal sequence residues at positions −15, −12, −7, and −4.

**Figure 8.**
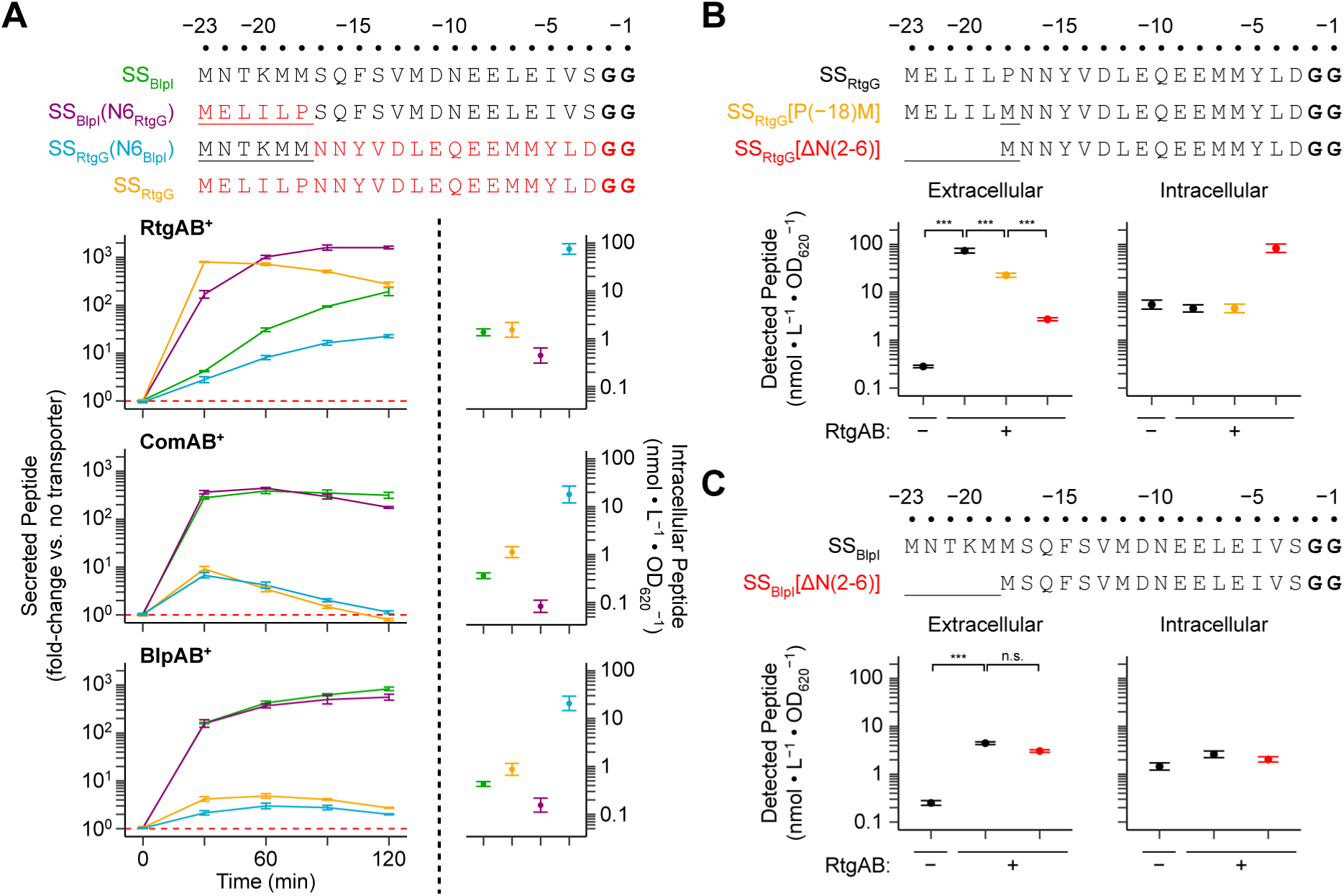
A unique motif found at the N-terminal ends of the *rtg* GG peptides promotes secretion by RtgAB. **(A)** R6 ComAB^−^/BlpAB^−^/RtgAB^−^ and single-transporter positive ComAB^+^, BlpAB^+^, RtgAB^+^ strains expressing BlpI or RtgG signal sequences with residue swaps at positions −22 to −18 (top; mutated positions are underlined) fused to BlpI-HiBiT cargo peptide were grown in CDM+ to OD_620_ 0.05 and treated with 200 ng/mL CSP, 200 ng/mL BlpC, and 20 nM RtgS_A_-C10. Samples were taken every 30 min and extracellular HiBiT signal was quantified (left). Data are presented as fold-change values relative to the ComAB^−^/BlpAB^−^/RtgAB^−^ control. Red, dashed line represents fold-change = 1 (no difference vs. the control). At the 120-min timepoint, intracellular peptide content was also quantified (right). Plots show mean ± S.E. of 3 independent experiments. **(B-C)** R6 ComAB^−^/BlpAB^−^/RtgAB^−^ and RtgAB^+^ strains expressing mutated SS_RtgG_ (B) or SS_BlpI_ (C) (top; mutated positions are underlined) fused to BlpI-HiBiT cargo peptide were grown and treated with pheromone as in panel A. Extracellular (left graphs) and intracellular (right graphs) HiBiT signals were quantified at 60 min post-treatment. Plots show mean ± S.E. of 3 independent experiments. Statistics: n.s. not significant, *** p < 0.001; ANOVA with Tukey’s HSD.

## Discussion

In this work, we have characterized the PCAT-encoding locus *rtg* and shown it is regulated by the RtgR/S system. RtgR/RtgS belongs to a family of regulatory systems found in streptococci that includes the Rgg/SHP and ComR/S systems (12). Rgg/SHP and ComR/S circuits can either act as cell density-dependent quorum sensing systems (12) or timing devices (40). Our data suggest RtgR/S behaves like the former (Fig. S9). A purely intracellular signaling pathway has been reported for XIP in *Streptococcus mutans* (14, 41). Such a pathway is unlikely to exist for RtgR/S, since *rtg* auto-induction requires both PptAB and Ami (Fig. S3A). While the RtgS pheromone is similar to the previously described SHP and ComS/XIP pheromones, it also differs from these other pheromone classes in important ways. RtgS lacks the conserved aspartate or glutamate residue characteristic of SHPs and is divergently transcribed from its regulator unlike ComS (12). However, RtgS does contain a Trp-Gly-Trp motif near the C-terminus which bears resemblance to the Trp-Trp motif found in some XIPs (12, 20). RtgR is phylogenetically closer to the ComRs than SHP-associated Rgg regulators but does not cluster with either group (12). Using a published list of Rgg regulators (12) we found two RtgR-like regulators associated with Trp-X-Trp (WxW) motif-containing pheromones: *SPD_1518* (*rgg1518*) from *S. pneumoniae* D39 and *SSA_2251* from *Streptococcus sanguinis* SK36 (Table S1). Expression analysis of the pheromone operon associated with *rgg1518* using PneumoExpress (42) revealed that the pheromone and genes *SPD_1513* to *SPD_1517* are specifically upregulated under the same conditions that result in upregulation of *rtg*. Therefore, the Rgg1518 system is likely functional. We propose that RtgR/S and other Rgg/WxW pheromone pairs constitute a distinct group of Rgg regulatory systems. We leave the work of characterizing the members of this group and the pathways they regulate to future studies.

We showed that in the RtgAB^+^ strain Sp9-BS68, the ability to activate the RtgR/S system confers a fitness advantage during competitive colonization of the nasopharynx. While 78% of strains are predicted to encode a functional RtgR and therefore can respond to pheromone, only 17% of strains are RtgAB^+^. Most RtgAB^−^ strains still encode at least one *rtg* GG peptide but have no obvious means with which to secrete them since they are not secreted by the other two PCATs commonly found in pneumococcus, ComAB and BlpAB. We have been unable to determine the function of the *rtg* GG peptides, but we speculate that they are bacteriocins. The reasons for this are that bacteriocin secretion is the most common function of PCATs, and five of the seven *rtg* GG peptide genes are always associated with downstream genes encoding hypothetical proteins that resemble bacteriocin immunity proteins (43). Regardless of the specific function of the *rtg* GG peptides, the fact that most RtgAB^−^ strains are still RtgR^+^ suggests that *rtg* retains a useful function that does not require secretion of these peptides. Further studies will be needed to determine the mechanism responsible for the RtgR/S-dependent competitive fitness advantage seen in colonization studies, the function of the *rtg* GG peptides, and the biological significance of active *rtg* loci with non-functional RtgAB.

The case of RtgAB and ComAB/BlpAB allowed us to study how two sets of PCATs which co-exist in the same strain preferentially secrete different sets of peptides through slight differences in substrate recognition. Unlike ComAB and BlpAB, RtgAB recognizes its substrates partially using a motif located at the N-terminal end of their signal sequences. This motif is located 18 residues away from the signal sequence cleavage site and is exclusively found in *rtg* GG peptides. Where data are available, previous studies of PCAT substrates have found that positions at the N-terminus located farther than 18 residues from the cleavage site are either dispensable for recognition by PCATs (28, 30) or can be missing entirely (33, 44, 45). As far as we are aware, RtgAB is unique among PCATs in recognizing a signal sequence motif located so distantly from the cleavage site. Future efforts will be directed toward identifying the specific nature of the interaction between the N-terminal motif and RtgAB and the exact signal sequence residues involved.

The insights into the sequence determinants of PCAT substrate selectivity gained here illuminates a relatively understudied aspect of this class of transporters. They will also be useful in guiding future efforts to predict substrates for ComAB, BlpAB, RtgAB, and other PCATs. Some GG peptides are found without a closely associated or co-regulated PCAT (18). In these cases, it would be helpful to have sequence-based approaches to assigning potential transporters to these “orphan” GG peptides. Moreover, for strains that encode multiple PCATs, predicting if GG peptides can be secreted by PCATs that are not necessarily closely associated can guide mechanistic studies that lead to new insights into function and regulation, such as with ComAB and BlpAB substrates in pneumococcus. Our work lays the groundwork for identifying signal sequence motifs of GG peptides that are important for transporter selectivity. The next step will be to study the corresponding sequence and structural motifs in PCATs that contribute to this selectivity. In addition to bacteriocins and quorum sensing (3), GG peptides have now been linked to biofilm formation, colonization of host niches, and dissemination during infection (18, 46). Ultimately, the ability to predict and rationalize PCAT-GG peptide pairings will advance our understanding of a broad range of biologically significant microbial processes.

## Materials and Methods

### Strains and growth conditions

All strains are derived from Sp9-BS68 (36), D39, or the R6 strain P654 (47) (referred to as PSD100 in reference) (see Table S2 and Supplemental Methods for details). Pneumococcus was grown in either filter-sterilized THY (Todd Hewitt broth + 0.5% yeast extract) or CDM+ (38) at 37°C. All media contained 5 µg/mL catalase. All CDM+ was supplemented with 0.5% (v/v) THY. Except where noted otherwise, pneumococcal cultures used for experiments were inoculated to OD_620_ 0.0015 from starter cultures grown in THY pH 7.4 to OD_620_ 0.275 and frozen at −80°C in 13% glycerol. Starter cultures were pelleted at 6000×*g*, 5 min, room temperature, and resuspended in the appropriate growth media for the experiment before being used for inoculation. Antibiotics were used at the following concentrations: chloramphenicol, 2 µg/mL; gentamicin, 200 µg/mL; kanamycin, 500 µg/mL; spectinomycin, 200 µg/mL; streptomycin, 100 µg/mL.

### Transformations

Transformation protocols were adapted from (48). See Supplemental Methods and Table S3 for details and primers used for constructing transforming DNA products. Unmarked chromosomal mutations were created via Janus (49), Sweet Janus (50), or Janus2 (see Supplemental Methods) exchange. Transformants were verified by Sanger sequencing.

### Luciferase reporter time course assays

For *com*/*blp* activation assays only, starter cultures were grown in THY pH 6.8 to OD_620_ 0.075 to prevent *com*/*blp* activation. Cells were grown in THY or CDM+ in a white, clear-bottom 96-well plate (Greiner Bio-One, 655098), 200 µL per well. For assays using firefly luciferase, the following concentrations of firefly luciferin (Thermo Fisher Scientific, 88294) were added to the media: 330 µM (single reporter and dual reporter, CDM+), 165 µM (dual reporter, THY). For assays using NanoLuc luciferase, the following concentrations of Nano-Glo substrate (Promega, N1121) were added to the media: 1:5000 (CDM+), 1:10000 (THY). The plate was incubated in a Synergy HTX plate reader, set to read absorbance at 620 nm and luminescence every 5 min. For single reporter assays, no filter was used for luminescence readings. For dual reporter assays, 450/50 band-pass and 610 long-pass filters were used to isolate NanoLuc and red firefly luciferase signals, respectively. For D39 strains only, the plate was shaken before readings were taken. Promoter activities were calculated from luminescence and absorbance readings as described in (25). For locus activation assays, timings of activation events were calculated as described in Supplemental Methods and compared using survival analysis. Differences between groups were assessed by log-rank tests using the FHtest package (v1.4) in R, and when appropriate the Holm correction was applied for multiple comparisons.

### RtgS dose-response assays

Cells expressing P*_rtgS1_*-*luc* reporters were grown in THY or CDM+ containing 330 µM firefly luciferin. At OD_620_ 0.02, cultures were aliquoted into a white, clear-bottom 96-well plate (Greiner Bio-One, 655098), 100 µL per well. Each well of the plate was prefilled with 100 µL sterile media containing 0.5% (v/v) DMSO, 330 µM firefly luciferin and appropriate concentrations of synthetic RtgS peptide (Genscript). The plate was then incubated in a Synergy HTX plate reader set to read absorbance at 620 nm and luminescence every 5 min. For D39 strains only, the plate was shaken before readings were taken. P*_rtgS1_* activity was calculated and the response was defined as the maximum observed P*_rtgS1_* activity within 60 min (Sp9-BS68) or 120 min (D39) of treatment. When applicable, curves were fit to a Hill model using the nls() function in R 3.5.1.

### Peptide secretion assays

Cells were inoculated from starter cultures to OD_620_ 0.005 and grown in CDM+. At OD_620_ 0.05, cells were treated with 200 ng/mL CSP1, 200 ng/mL BlpC_R6_, and 20 nM RtgS_A_-C10. Samples were taken for HiBiT quantification at appropriate time points. For native BlpI-HiBiT assays only, clarified supernatants were obtained after centrifugation at 6000×*g*, 5 min, 4°C. For all other assays, cells were retained in the samples. HiBiT signal was quantified by mixing samples with HiBiT Extracellular Detection Reagent (Promega, N2421) at a 1:1 ratio and reading luminescence with a Synergy HTX plate reader. Samples were also taken for quantification of intracellular peptide; for endpoint assays, they were taken concurrently with the extracellular samples, and for time-course assays, they were taken at the last time point. Extracellular peptide was removed from these samples by proteinase K digestion, then the cells were lysed and HiBiT signal was quantified as above. Standards consisting of synthetic L10-HiBiT peptide (25) mixed with samples of a non-HiBiT expressing strain were used to generate standard curves to use for calculating HiBiT-tagged peptide concentrations in experimental samples. See Supplemental Methods for more details. Differences between groups were assessed by ANOVA, using the emmeans package (v1.2.3) in R.

### Genomic analysis of rtg

Analysis of *rtg* was performed using the assembled genomes of the Massachusetts isolate collection (BioProject Accession: PRJEB2632). See Supplemental Methods for details.

### Mouse colonization assays

Mouse colonization was performed as described in (25). Briefly, dual or single-strain mixtures of Sp9-BS68 were inoculated into the nasopharynx of un-anaesthetized 5- to 7-week-old female BALB/c mice (Taconic). Mice were euthanized with CO_2_ overdose after 72 hours and nasopharyngeal colonization was sampled by nasal wash. See Supplemental Methods for IACUC approval and details on how colonization density and competitive indices were calculated. Differences in colonization densities and competitive indices between groups were evaluated by the Mann-Whitney (2 groups) and Kruskal-Wallis (>2 groups) tests using the wilcox.test() and kruskal.test() functions in R 3.5.1.

## Acknowledgments

This work was supported by funding from the National Institutes of Health T32 AI007528 (CW) and R56 AI101285 (SD).

## Supplemental Methods

### DNA manipulation

All PCR reactions for downstream Gibson assembly, transformation, or sequencing applications were performed using Phusion polymerase (NEB, M0530). Other PCR reactions were performed using Taq polymerase (NEB, M0273). For PCR reactions in which cells were used as template, the cells were obtained from starter cultures resuspended in sterile water and added to the PCR reaction at a 1:20 (Phusion) or 1:10 (Taq) dilution. For PCR reactions in which crude Gibson assembly product was used as template, the crude Gibson assembly product was added to the PCR reaction at a 1:100 dilution. Gibson assembly was performed using NEB HiFi DNA Assembly master mix (NEB, E2621). Primers were designed with the aid of primer3 (1, 2) and synthesized by IDT.

### Construction of the Janus+ cassette

Two PCR products, amplified using primer pairs CW105/CW357 on Janus cassette (3) template, and CW358/CW106 on Sweet Janus+ cassette (4) template, were ligated via Gibson assembly. The resulting 1402-bp product formed the Janus+ cassette, which contains the *aphA-3* gene (kanamycin resistance) driven by the pneumococcal *amiA* promoter, followed by the *rpsL+* gene (dominant-negative streptomycin sensitivity) driven by the pneumococcal *rplM* promoter. The version of *rpsL+* included in Janus+ has the same five silent substitutions as the version in Sweet Janus+. These substitutions and the increased expression from the *rplM* promoter collectively reduce the rate of spontaneous reversion to a streptomycin resistant phenotype in strains carrying the Janus+ cassette compared to the original Janus cassette.

### Construction of the Janus2 cassette

A gBlocks DNA fragment was synthesized by IDT containing the gentamicin resistance gene *aacC1* from pPEPY-PF6-*lacI* (5). This template was PCR amplified using primers CW407/CW406 and ligated via Gibson assembly with another PCR product amplified using primers CW105/CW375 from Sweet Janus+ cassette (4) template. The resulting 2069-bp product formed the Janus2 cassette, which contains the *sacB* and *aacC1* genes in a single operon driven by pneumococcal *amiA* promoter.

### Allelic exchange using the Janus2 cassette

The Janus2 cassette confers gentamicin resistance (200 µg/mL) and sensitivity to 10% sucrose to pneumococcal strains into which it is inserted. This allows it to be used as a counterselectable marker in the same vein as the Janus (3) and Sweet Janus cassettes (6) on which it is based. Initial selection after transforming the Janus2 cassette into pneumococcus is performed using gentamicin. Afterwards, the cassette can be exchanged with an arbitrary, markerless DNA fragment. Growth in the presence of 10% sucrose selects for transformants that have successfully exchanged the Janus2 cassette. Because the negative selection step for Janus2 does not require a streptomycin resistant copy of *rpsL* in the chromosome (or any other specific feature), Janus2 theoretically can be used in any pneumococcal strain “as is”. Also, the Janus2 cassette and the original Janus cassette (or Janus+ cassette) use orthogonal selection agents. Therefore, Janus2 can be inserted into a strain already harboring Janus, and vice versa. Exchange of the two cassettes can then be performed in either order. Alternatively, insertion and exchange of the cassettes can each be performed in a single transformation, halving the number of transformation steps required to perform allelic exchange at two unlinked loci.

### Construction of the Sp9-BS68 constitutive luciferase reporter

The Sp9-BS68 PF6-*luc* reporter in which the firefly luciferase gene (*luc*) is expressed from the highly active PF6 promoter (5) was created as follows. First, a PCR product (Sp9-BS68-CEP-Janus2-*luc*) of the Janus2 cassette and *luc* inserted into the CEP site (7) between *treR* and *amiF* was amplified using primers CW303/CW295 following Gibson assembly of four PCR products amplified using primer pairs CW456/CW454 on Sp9-BS68 template, CW105/CW406 on Janus2 cassette template, CW457/CW458 on P1666 (4) template, and CW343/CW294 on a GenParts fragment (Genscript) containing the *thrC* terminator from *E. coli* (8), the *tufA* terminator from pneumococcus (9), and approximately 500 bp of flanking sequence including *treR*. The assembled product contains a deletion of the region from n.t. +639 of the gene *CGSSp9BS68_00992* to n.t. −466 of the gene *CGSSp9BS68_00972*, into which are inserted the Janus2 cassette followed immediately by *luc*, oriented in the same direction as *amiF*, then a 50-n.t. spacer and finally the two terminators. The deletion removes an ABC transporter operon predicted to encode a non-functional product due to disruptions of the putative substrate-binding protein and permease genes. Second, a PCR product (Sp9-BS68-CEP-PF6-*luc*) of the luciferase reporter was amplified using primers CW463/CW188 following Gibson assembly of three PCR products amplified using primer pairs CW464/CW455 on Sp9-BS68 template, CW380/CW190 on a GenParts fragment (Genscript) containing the *hisI* and *rpsI* terminators from *E. coli* (8), the PF6 promoter, and the first 103 n.t. of *luc*, and CW191/CW189 on P1666 template. The Sp9-BS68-CEP-Janus2-*luc* PCR product was transformed into Sp9-BS68 to create strain P2769 and then the Janus2 cassette was exchanged with the Sp9-BS68-CEP-PF6-*luc* PCR product to create strain P2772.

### Construction of Sp9-BS68 rtg luciferase reporters

The Sp9-BS68 P*_rtgA_*-*luc* reporter in which the firefly luciferase gene (*luc*) is inserted in place of *rtgA* following an ectopic copy of the *rtgS1*/*rtgA* promoter was created as follows. A PCR product (Sp9-BS68-CEP-P*_rtgA_*-*luc*) was amplified using primers CW463/CW188 following Gibson assembly of three PCR products amplified using primer pairs CW464/CW455 on Sp9-BS68 template, CW380/CW190 on a GenParts fragment (Genscript) containing the *hisI* and *rpsI* terminators from *E. coli*, the region from n.t. −429 to n.t. −1 of *rtgA*, and the first 103 n.t. of *luc*, and CW191/CW189 on P1666 template. Then, the Janus2 cassette in P2769 was exchanged with the Sp9-BS68-CEP-P*_rtgA_*-*luc* PCR product to create strain P2775. While the ectopic promoter region of *rtgA* in front of *luc* in P2775 contains a copy of *rtgS1*, this copy is disrupted by a frameshift mutation. Therefore, P2775 still only has one functional copy of *rtgS1*.

The Sp9-BS68 P*_rtgS1_*-*luc* reporter in which the firefly luciferase gene (*luc*) is inserted in place of *rtgS1* following an ectopic copy of the *rtgS1* promoter was created as follows. A PCR product (Sp9-BS68-CEP-P*_rtgS1_*-*luc*) was amplified using primers CW463/CW188 following Gibson assembly of two PCR products amplified using primer pairs CW464/CW494 and CW158/CW189 on P2775 template. Then, the Janus2 cassette in P2769 was exchanged with the Sp9-BS68-CEP-P*_rtgS1_*-*luc* PCR product to create strain P2792.

### Construction of D39 rtg luciferase reporter

The D39 P*_rtgS1_*-*luc* reporter in which the firefly luciferase gene (*luc*) is inserted in place of *rtgS1* following an ectopic copy of the *rtgS1* promoter was created as follows. First, a PCR product (D39-CEP-Janus2-*luc*) of the Janus2 cassette and *luc* inserted into the CEP site between *treR* and *amiF* was amplified using primers CW463/CW293 following Gibson assembly of three PCR products amplified using primer pairs CW464/CW471 on P2055 (4) template, CW105/CW474 on P2769 template, and CW481/CW292 on P2055 template. The assembled product contains a deletion of the region from n.t. −101 to n.t. +1168 of the gene *SPD_1666*, replaced by the insertion from P2769. The deletion removes a degenerate transposon. Second, a PCR product (Sp9-BS68-CEP-P*_rtgS1_*-*luc*) containing the luciferase reporter was amplified using primers CW463/CW188 following Gibson assembly of three PCR products amplified using primer pairs CW464/CW482 on P2055 template, CW380/CW494 on P2775 template, and CW158/CW189 on P1666 template. The D39-CEP-Janus2-*luc* PCR product was transformed into P2055 to create strain P2779 and then the Janus2 cassette was exchanged with the D39-CEP-P*_rtgS1_*-*luc* PCR product to create strain P2790.

### Construction of Sp9-BS68 rtgR and rtgS1 deletion mutants

The Sp9-BS68 Δ*rtgR* strain was created as follows. First, a PCR product (Sp9-BS68-*rtgR*-Janus2) was amplified using primers CW487/CW486 following Gibson assembly of three PCR products amplified using primer pairs CW488/CW490 on Sp9-BS68 template, CW105/CW406 on Janus2 cassette template, and CW489/CW485 on Sp9-BS68 template. The assembled product contains the Janus2 cassette inserted in place of *rtgR*. Second, a PCR product (Sp9-BS68-Δ*rtgR*) was amplified using primers CW487/CW486 following Gibson assembly of two PCR products amplified using primer pairs CW488/CW491 and CW492/CW485 on Sp9-BS68 template. The Sp9-BS68-*rtgR*-Janus2 PCR product was transformed into P2792 and then the Janus2 cassette was exchanged with the Sp9-BS68-Δ*rtgR* PCR product to create strain P2802.

The Sp9-BS68 Δ*rtgS1* strain was created as follows. First, a PCR product (Sp9-BS68-*rtgR*-*rtgS1*-Janus2) was amplified using primers CW498/CW486 following Gibson assembly of three PCR products amplified using primer pairs CW499/CW523 on Sp9-BS68 template, CW105/CW406 on Janus2 cassette template, and CW489/CW485 on Sp9-BS68 template. The assembled product contains the Janus2 cassette inserted in place of *rtgR* and *rtgS1*. Second, a PCR product (Sp9-BS68-Δ*rtgS1*) was amplified using primers CW498/CW486 following Gibson assembly of three PCR products amplified using primer pairs CW488/CW503 on Sp9-BS68 template, CW502/CW484 on a gBlocks fragment (IDT) containing an in-frame deletion of n.t. +31 to n.t. +84 of *rtgS1*, and CW483/CW485 on Sp9-BS68 template. The Sp9-BS68-*rtgR*-*rtgS1*-Janus2 PCR product was transformed into P2792 to create strain P2798 and then the Janus2 cassette was exchanged with the Sp9-BS68-Δ*rtgS1* PCR product to create strain P2804.

The Sp9-BS68 Δ*rtgR*Δ*rtgS1* strain was created as follows. A PCR product (Sp9-BS68-Δ*rtgR*Δ*rtgS1*) was amplified using primers CW498/CW486 following Gibson assembly of two PCR products amplified using primer pairs CW488/CW484 on P2804 template and CW483/CW485 on P2802 template. Then, the Janus2 cassette in P2798 was exchanged with the Sp9-BS68-Δ*rtgR*Δ*rtgS1* PCR product to create strain P2811.

### Construction of D39 rtgS1 and rtgS2 deletion mutants

The D39 Δ*rtgS1* strain was created as follows. First, a PCR product (D39-*rtgS1*-Janus2) was amplified using primers CW556/CW498 following Gibson assembly of three PCR products amplified using primer pairs CW496/CW500 on P2055 template, CW105/CW406 on Janus2 cassette template, and CW501/CW499 on P2055 template. Second, a PCR product (D39-Δ*rtgS1*) was amplified using primers CW556/CW498 following Gibson assembly of three PCR products amplified using primer pairs CW496/CW483 on P2055 template, CW484/CW502 on P2804 template, and CW503/CW499 on P2055 template. The D39-*rtgS1*-Janus2 PCR product was transformed into P2790 to create strain P2851 and then the Janus2 cassette was exchanged with the D39-Δ*rtgS1* PCR product to create strain P2859.

The D39 Δ*rtgS2* strain was created as follows. First, a PCR product (D39-*rtgS2*-Janus+) was amplified using primers CW513/CW518 following Gibson assembly of three PCR products amplified using primer pairs CW548/CW562 on P2055 template, CW105/CW106 on Janus+ cassette template, and CW561/CW519 on P2055 template. Second, a PCR product (D39-Δ*rtgS2*) was amplified using primers CW513/CW518 following Gibson assembly of three PCR products amplified using primer pairs CW548/CW483 on P2055 template, CW484/CW520 on a gBlocks fragment (IDT) containing an in-frame deletion of n.t. +31 to n.t. +84 of *rtgS2*, and CW521/CW519 on P2055 template. Third, a PCR product (D39-*rtgS1*) was amplified using primers CW556/CW498 on P2055 template. The D39-*rtgS2*-Janus+ PCR product was transformed into P2851 to create strain P2896. Then, the Janus+ cassette was exchanged with the D39-Δ*rtgS2* PCR product to create strain P2902. Finally, the Janus2 cassette was exchanged with the D39-*rtgS1* PCR product to create strain P2908.

The D39 Δ*rtgS1*Δ*rtgS2* strain was created as follows. The Janus2 cassette in strain P2902 was exchanged with the D39-Δ*rtgS1* PCR product to create strain P2910.

### Construction of Sp9-BS68 ami and ppt deletion mutants

The Sp9-BS68 Δ*amiCD* strain was created as follows. First, a PCR product (Sp9-BS68-*amiCD*-Janus2) was amplified using primers CW616/CW607 following Gibson assembly of three PCR products amplified using primer pairs CW617/CW614 on Sp9-BS68 template, CW105/CW406 on Janus2 cassette template, and CW611/CW606 on Sp9-BS68 template. Second, a PCR product (Sp9-BS68-Δ*amiCD*) was amplified using primers CW616/CW607 following Gibson assembly of two PCR products amplified using primer pairs CW617/CW613 and CW612/CW606 on Sp9-BS68 template. The Sp9-BS68-*amiCD*-Janus2 PCR product was transformed into P2792, and then the Janus2 cassette was exchanged with the Sp9-BS68-Δ*amiCD* PCR product to create strain P3075.

The Sp9-BS68 Δ*pptAB* strain was created as follows. First, a PCR product (Sp9-BS68-*pptAB*-Janus2) was amplified using primers CW619/CW627 following Gibson assembly of three PCR products amplified using primer pairs CW618/CW622 on Sp9-BS68 template, CW105/CW406 on Janus2 cassette template, and CW625/CW628 on Sp9-BS68 template. Second, a PCR product (Sp9-BS68-Δ*amiCD*) was amplified using primers CW619/CW627 following Gibson assembly of two PCR products amplified using primer pairs CW618/CW623 and CW624/CW628 on Sp9-BS68 template. The Sp9-BS68-*pptAB*-Janus2 PCR product was transformed into P2792, and then the Janus2 cassette was exchanged with the Sp9-BS68-Δ*pptAB* PCR product to create strain P3077.

### Construction of Sp9-BS68 rtgS1 promoter mutation reporters

The Sp9-BS68 P*_rtgS1_*(P2)-*luc* reporter strain was created as follows. First, the Sp9-BS68-CEP-Janus2-*luc* PCR product was amplified using primers CW463/CW468 on P2769 template. Second, a PCR product (Sp9-BS68-CEP-P*_rtgS1_*(P2)-*luc*) was amplified using primers CW463/CW188 following Gibson assembly of two PCR products amplified using primer pairs CW464/CW570 and CW629/CW189 on P2792 template. The Sp9-BS68-CEP-Janus2-*luc* PCR product was transformed into strain P2804 to create strain P3080, and then the Janus2 cassette was exchanged with the Sp9-BS68-CEP-P*_rtgS1_*(P2)-*luc* PCR product to create strain P3100.

The Sp9-BS68 P*_rtgS1_*(P2)-*luc* reporter strains with promoter sequence mutations were created as follows. The mut1, mut2, mut3, and mut4 PCR products were amplified using primers CW463/CW188 following Gibson assembly of the following pairs of PCR products, all amplified from P3100 template: mut1, CW464/CW632 and CW633/CW189; mut2, CW464/CW630 and CW631/CW189; mut3, CW464/CW634 and CW635/CW189; mut4, CW464/CW636 and CW637/CW189. The Janus2 cassette in strain P3080 was then exchanged with mut1, mut2, mut3, and mut4 to create strains P3123, P3125, P3127, and P3129, respectively.

### Construction of D39 com/blp luciferase reporters with repaired rtgA (RtgAB^+^)

First, a PCR product (D39-*rtgAXB*-SJanus+) was amplified using primers CW556/CW557 following Gibson assembly of three PCR products amplified using primer pairs CW496/CW537 on D39 template, CW105/CW106 on Sweet Janus+ cassette template, and CW509/CW559 on D39 template. Second, a PCR product (D39-*rtgAXB_Sp9-BS68_*) was amplified using primers CW556/CW557 following Gibson assembly of three PCR products amplified using primer pairs CW496/CW502 on D39 template, CW503/CW511 on Sp9-BS68 template, and CW522/CW559 on D39 template. The D39-*rtgAXB*-SJanus+ PCR product was transformed into strains P2665, P2666, P2668, and P2670 and then the Sweet Janus+ cassettes were exchanged with the D39-*rtgAXB_Sp9-BS68_* PCR product to create strains P2838, P2840, P2842, and P2844, respectively.

### Construction of Sp9-BS68 strains for mouse colonization assays

The spectinomycin-resistant Sp9-BS68 strains were created as follows. A PCR product (Sp9-BS68-CEP-spcR) was amplified using primers CW463/CW468 following Gibson assembly of three PCR products amplified using primer pairs CW464/CW570 on P2792 template, CW571/CW572 on pE81 template, and CW343/CW292 on P2792 template. The Sp9-BS68-CEP-spcR PCR product was transformed into strains P2792 and P2811 to create strains P3001 and P3003, respectively.

The gentamicin-resistant Sp9-BS68 strain was created as follows. A PCR product (Sp9-BS68-CEP-genR) was amplified using primers CW463/CW468 following Gibson assembly of three PCR products amplified using primer pairs CW464/CW600 on P3001 template, CW598/CW599 on Janus2 cassette template, and CW343/CW292 on P3001 template. The Sp9-BS68-CEP-genR PCR product was transformed into strain P2811 to create strain P3025.

Mouse-passaged versions of the above strains were created as follows. P3001, P3003, and P3025 were inoculated into 5-7-week-old female BALB/c mice using the same protocol as that used for the single-strain colonization assays. After 24 hours, the colonizers were recovered via nasal wash using the same protocol as that used for the colonization assays. Nasal washes were plated on spectinomycin- or gentamicin-containing TSA plates supplemented with 5 µg/mL catalase and incubated overnight at 37°C with 5% CO_2_. From each of the P3001, P3003, and P3025-inoculated mice, eight individual colonies were pooled to create stocks of strains P3035, P3037, and P3039, respectively. These mouse-passaged strains were then used for the mouse colonization assays.

### Construction of rtg autoinducing-deficient tagged BlpI expressing R6 strains

First, a PCR product (R6-*rtgS1*-*rtgD2*-SJanus+) was amplified using primers CW556/CW550 following Gibson assembly of three PCR products amplified using primer pairs CW496/CW500 on R6 template, CW105/CW106 on Sweet Janus+ cassette template, and CW538/CW551 on R6 template. Second, a PCR product (R6-Δ*rtgS1*-*rtgAXB*) was amplified using primers CW556/CW550 following Gibson assembly of two PCR products amplified using primers CW556/CW565 on P2859 template, and CW542/CW551 on R6 template. Third, a PCR product (R6-Δ*rtgS1*-*rtgAXB_Sp9-BS68_*) was amplified using primers CW556/CW550 following Gibson assembly of three PCR products amplified using primers CW556/CW483 on R6 template, CW484/CW541 on P2804 template, and CW542/CW551 on R6 template. The R6-*rtgS1*-*rtgD2*-SJanus+ PCR product was transformed into strains P2538, P2565, P2567, and P2569 to create strains P2865, P2867, P2869, and P2871, respectively. Then, the Sweet Janus+ cassettes in P2865, P2867, P2869, and P2871 were exchanged for the R6-Δ*rtgS1*-*rtgAXB* PCR product to create strains P2878, P2880, P2884, and P2888, respectively. Finally, the Sweet Janus+ cassette in strain P2867 was exchanged for the R6-Δ*rtgS1*-*rtgAXB_Sp9-BS68_* PCR product to create strain P2882.

### Construction of RtgC-HiBiT and RtgG-HiBiT expressing R6 strains

The R6 strains expressing RtgC-HiBiT were created as follows. First, the pE57 insertion and *blpI_P133_*-*HiBiT* genes were removed from P2880, P2882, P2884, and P2888. A PCR product (P654-*blpA*-*pncS*-Janus+) was amplified using primers CW271/CW219 following Gibson assembly of three PCR products amplified using primer pairs CW272/CW270 on P654 (10) template, CW105/CW106 on Janus+ cassette template, and CW238/CW234 on P654 template. Another PCR product (P654-ΔpE57) was amplified using primers CW585/CW219 following Gibson assembly of two PCR products amplified using primer pairs CW584/CW586 and CW587/CW234 on P654 template. The P654-*blpA*-*pncS*-Janus+ PCR product was transformed into strains P2880, P2882, P2884, and P2888 and then exchanged with the P654-ΔpE57 PCR product to create strains P2934, P2936, P2938, and P2940, respectively. Second, *rtgC-HiBiT* was inserted downstream of *rtgB* in these strains. A PCR product (P2880-*rtgC*-*rtgD2*-SJanus+) was amplified using primers CW451/CW550 following Gibson assembly of three PCR products amplified using primer pairs CW531/CW534 on R6 template, CW105/CW106 on Sweet Janus+ cassette template, and CW538/CW551 on R6 template. Another PCR product (P2882-*rtgC*-*rtgD2*-SJanus+) was amplified using primers CW451/CW550 following Gibson assembly of three PCR products amplified using primer pairs CW543/CW546 on Sp9-BS68 template, CW105/CW106 on Sweet Janus+ cassette template, and CW538/CW551 on R6 template. A third PCR product (P2880-*rtgC-HiBiT*) was amplified using primers CW451/CW550 following Gibson assembly of three PCR products amplified using primer pairs CW531/CW506 on R6 template, CW592/CW593 on a gBlocks fragment (IDT) containing *rtgC_R6_* with an insertion of sequence encoding HiBiT tag preceded by a 10-residue linker (ggtggtggaggttcaggaggtggaggttctgtttctggttggcgtctttttaaaaaaatttca) followed by *rtgD1_R6_*, and CW594/CW551 on R6 template. A final PCR product (P2882-*rtgC-HiBiT*) was amplified using primers CW451/CW550 following Gibson assembly of three PCR products amplified using primer pairs CW543/CW546 on a template containing *rtgAXB* from Sp9-BS68 inserted in place of *rtgAXB* in the D39 *rtg* locus, CW592/CW593 on the *rtgC-HiBiT* gBlocks fragment from above, and CW594/CW551 on R6 template. The P2880-*rtgC*-*rtgD2*-SJanus+ PCR product was transformed into strains P2934, P2938, and P2940 to create strains P2945, P2949, and P2951, respectively. The P2882-*rtgC*-*rtgD2*-SJanus+ PCR product was transformed into strain P2936 to create strain P2947. Then, the Sweet Janus+ cassettes in strains P2945, P2949, and P2951 were exchanged for the P2880-*rtgC-HiBiT* PCR product to create strains P2959, P2963, and P2965, respectively. Finally, the Sweet Janus+ cassette in strain P2947 was exchanged for the P2882-*rtgC-HiBiT* PCR product to create strain P2961.

The R6 strains expressing RtgG-HiBiT were created as follows. A PCR product (P2880-*rtgG-HiBiT*) was amplified using primers CW451/CW550 following Gibson assembly of three PCR products amplified using primer pairs CW531/CW595 on R6 template, CW522/CW593 on a gBlocks fragment (IDT) containing *rtgG_Sp9-BS68_* with an insertion of sequence encoding HiBiT tag preceded by a 10-residue linker (ggtggtggaggttcaggaggtggaggttctgtttctggttggcgtctttttaaaaaaatttca) followed by *rtgH_Sp9-BS68_*, and CW594/CW551 on R6 template. Another PCR product (P2882-*rtgG-HiBiT*) was amplified using primers CW451/CW550 following Gibson assembly of three PCR products amplified using primer pairs CW543/CW511 on P2882 template, CW522/CW593 on the *rtgG-HiBiT* gBlocks fragment from above, and CW594/CW551 on R6 template. The Sweet Janus+ cassettes in strains P2945, P2949, and P2951 were exchanged for the P2880-*rtgG-HiBiT* PCR product to create strains P2967, P2971, and P2973, respectively. The Sweet Janus+ cassette in strain P2947 was exchanged for the P2882-*rtgG-HiBiT* PCR product to create strain P2969.

### Construction of the R6 strain with rtgA_ATG>ATT_ expressing RtgG-HiBiT

A PCR product (P2882-*rtgA*-Janus2) was amplified using primers CW556/CW659 following Gibson assembly of three PCR products amplified using primer pairs CW496/CW537 on P2882 template, CW105/CW406 on Janus2 cassette template, and CW657/CW660 on P2882 template. Another PCR product (P2882-*rtgA_ATG>ATT_*) was amplified using primers CW556/CW659 following Gibson assembly of three PCR products amplified using primer pairs CW496/CW502 on P2882 template, CW503/CW446 on R6 template, and CW447/CW660 on P2882 template. The P2882-*rtgA*-Janus2 PCR product was transformed into P2969, and then the Janus2 cassette was exchanged with the P2882-*rtgA_ATG>ATT_* PCR product to create strain P3162.

### Construction of R6 strains expressing BlpI-HiBiT from downstream of rtgB

A PCR product (P2880-*blpI-HiBiT*) was amplified using primers CW451/CW550 following Gibson assembly of three PCR products amplified using primer pairs CW531/CW595 on R6 template, CW522/CW596 on a gBlocks fragment (IDT) containing *blpI-HiBiT*, and CW597/CW551 on P2967 template. Another PCR product (P2882-*blpI-HiBiT*) was amplified using primers CW451/CW550 following Gibson assembly of three PCR products amplified using primer pairs CW543/CW511 on P2882 template, CW522/CW596 on the *blpI-HiBiT* gBlocks fragment from above, and CW597/CW551 on P2967 template. The Sweet Janus+ cassettes in strains P2945, P2949, and P2951 were exchanged for the P2880-*blpI-HiBiT* PCR product to create strains P3027, P3031, and P3033, respectively. The Sweet Janus+ cassette in strain P2947 was exchanged for the P2882-*blpI-HiBiT* PCR product to create strain P3029.

### Construction of R6 strains expressing signal sequence-swapped RtgG-HiBiT and BlpI-HiBiT

The R6 strains expressing SS_RtgG_-BlpI-HiBiT were created as follows. A PCR product (P2880-*SS_rtgG_-blpI-HiBiT*) was amplified using primers CW451/CW550 following Gibson assembly of three PCR products amplified using primer pairs CW531/CW595 on R6 template, CW522/CW596 on a gBlocks fragment (IDT) containing *SS_rtgG_-blpI-HiBiT*, and CW597/CW551 on P2967 template. Another PCR product (P2882-*SS_rtgG_-blpI*-*HiBiT*) was amplified using primers CW451/CW550 following Gibson assembly of three PCR products amplified using primer pairs CW543/CW511 on P2882 template, CW522/CW596 on the *SS_rtgG_-blpI-HiBiT* gBlocks fragment from above, and CW597/CW551 on P2967 template. The Sweet Janus+ cassettes in strains P2945, P2949, and P2951 were exchanged for the P2880-*SS_rtgG_-blpI-HiBiT* PCR product to create strains P2981, P2985, and P2987, respectively. The Sweet Janus+ cassette in strain P2947 was exchanged for the P2882-*SS_rtgG_-blpI-HiBiT* PCR product to create strain P2983.

The R6 strains expressing SS_BlpI_-RtgG-HiBiT were created as follows. A PCR product (P2880-*SS_blpI_-rtgG-HiBiT*) was amplified using primers CW451/CW550 following Gibson assembly of three PCR products amplified using primer pairs CW531/CW595 on R6 template, CW522/CW596 on a gBlocks fragment (IDT) containing *SS_blpI_-rtgG-HiBiT*, and CW597/CW551 on P2967 template. Another PCR product (P2882-*SS_blpI_-rtgG*-*HiBiT*) was amplified using primers CW451/CW550 following Gibson assembly of three PCR products amplified using primer pairs CW543/CW511 on P2882 template, CW522/CW596 on the *SS_blpI_-rtgG-HiBiT* gBlocks fragment from above, and CW597/CW551 on P2967 template. The Sweet Janus+ cassettes in strains P2945, P2949, and P2951 were exchanged for the P2880-*SS_blpI_-rtgG-HiBiT* PCR product to create strains P2993, P2997, and P2999, respectively. The Sweet Janus+ cassette in strain P2947 was exchanged for the P2882-*SS_blpI_-rtgG-HiBiT* PCR product to create strain P2995.

### Construction of R6 strains expressing HiBiT-tagged BlpI cargo peptide fused to various signal sequences

The R6 strains expressing SS_RtgG_(F/M/L/V)-BlpI-HiBiT were created as follows. A PCR product (P2880-*SS_rtgG_(F/M/L/V)-blpI-HiBiT*) was amplified using primers CW451/CW550 following Gibson assembly of three PCR products amplified using primer pairs CW531/CW595 on R6 template, CW522/CW596 on a gBlocks fragment (IDT) containing *SS_rtgG_(F/M/L/V)-blpI-HiBiT*, and CW597/CW551 on P2967 template. Another PCR product (P2882-*SS_rtgG_(F/M/L/V)-blpI-HiBiT*) was amplified using primers CW451/CW550 following Gibson assembly of three PCR products amplified using primer pairs CW543/CW511 on P2882 template, CW522/CW596 on the *SS_rtgG_(F/M/L/V)-blpI-HiBiT* gBlocks fragment from above, and CW597/CW551 on P2967 template. The Sweet Janus+ cassettes in strains P2945, P2949, and P2951 were exchanged for the P2880-*SS_rtgG_(F/M/L/V)-blpI-HiBiT* PCR product to create strains P3009, P3013, and P3015, respectively. The Sweet Janus+ cassette in strain P2947 was exchanged for the P2882-*SS_rtgG_(F/M/L/V)-blpI-HiBiT* PCR product to create strain P3011.

The R6 strains expressing SS_BlpI_(Y/L/M/L)-BlpI-HiBiT were created as follows. A PCR product (P2880-*SS_blpI_(Y/L/M/L)-blpI-HiBiT*) was amplified using primers CW451/CW550 following Gibson assembly of three PCR products amplified using primer pairs CW531/CW595 on R6 template, CW522/CW596 on a gBlocks fragment (IDT) containing *SS_blpI_(Y/L/M/L)-blpI-HiBiT*, and CW597/CW551 on P2967 template. Another PCR product (P2882-*SS_blpI_(Y/L/M/L)-blpI-HiBiT* was amplified using primers CW451/CW550 following Gibson assembly of three PCR products amplified using primer pairs CW543/CW511 on P2882 template, CW522/CW596 on the *SS_blpI_(Y/L/M/L)-blpI-HiBiT* gBlocks fragment from above, and CW597/CW551 on P2967 template. The Sweet Janus+ cassettes in strains P2945, P2949, and P2951 were exchanged for the P2880-*SS_blpI_(Y/L/M/L)-blpI-HiBiT* PCR product to create strains P3017, P3021, and P3023, respectively. The Sweet Janus+ cassette in strain P2947 was exchanged for the P2882-*SS_blpI_(Y/L/M/L)-blpI-HiBiT* PCR product to create strain P3019.

The R6 strains expressing SS_RtgG_(A/A/A/A)-BlpI-HiBiT were created as follows. A PCR product (P2880-*SS_rtgG_(A/A/A/A)-blpI-HiBiT*) was amplified using primers CW451/CW550 following Gibson assembly of three PCR products amplified using primer pairs CW531/CW595 on R6 template, CW522/CW596 on a gBlocks fragment (IDT) containing *SS_rtgG_(A/A/A/A)-blpI-HiBiT*, and CW597/CW551 on P2967 template. Another PCR product (P2882-*SS_rtgG_(A/A/A/A)-blpI-HiBiT*) was amplified using primers CW451/CW550 following Gibson assembly of three PCR products amplified using primer pairs CW543/CW511 on P2882 template, CW522/CW596 on the *SS_rtgG_(A/A/A/A)-blpI-HiBiT* gBlocks fragment from above, and CW597/CW551 on P2967 template. The Sweet Janus+ cassettes in strain P2945 was exchanged for the P2880-*SS_rtgG_(A/A/A/A)-blpI-HiBiT* PCR product to create strain P3111. The Sweet Janus+ cassette in strain P2947 was exchanged for the P2882-*SS_rtgG_(A/A/A/A)-blpI-HiBiT* PCR product to create strain P3113.

The R6 strains expressing SS_BlpI_(A/A/A/A)-BlpI-HiBiT were created as follows. A PCR product (P2880-*SS_blpI_(A/A/A/A)-blpI-HiBiT*) was amplified using primers CW451/CW550 following Gibson assembly of three PCR products amplified using primer pairs CW531/CW595 on R6 template, CW522/CW596 on a gBlocks fragment (IDT) containing *SS_blpI_(A/A/A/A)-blpI-HiBiT*, and CW597/CW551 on P2967 template. Then, the Sweet Janus+ cassettes in strains P2945, P2949, and P2951 were exchanged for the P2880-*SS_blpI_(A/A/A/A)-blpI-HiBiT* PCR product to create strains P3142, P3144, and P3146, respectively.

The R6 strains expressing SS_RtgG_(N6_BlpI_)-BlpI-HiBiT were created as follows. A PCR product (P2880-*SS_rtgG_(N6_blpI_)-blpI-HiBiT*) was amplified using primers CW451/CW550 following Gibson assembly of three PCR products amplified using primer pairs CW531/CW595 on R6 template, CW522/CW596 on a gBlocks fragment (IDT) containing *SS_rtgG_(N6_blpI_)-blpI-HiBiT*, and CW597/CW551 on P2967 template. Another PCR product (P2882-*SS_rtgG_(N6_blpI_)-blpI-HiBiT*) was amplified using primers CW451/CW550 following Gibson assembly of three PCR products amplified using primer pairs CW543/CW511 on P2882 template, CW522/CW596 on the *SS_rtgG_(N6_blpI_)*-*blpI-HiBiT* gBlocks fragment from above, and CW597/CW551 on P2967 template. The Sweet Janus+ cassettes in strains P2945, P2949, and P2951 were exchanged for the P2880-*SS_rtgG_(N6_blpI_)-blpI-HiBiT* PCR product to create strains P3055, P3059, and P3061, respectively. The Sweet Janus+ cassette in strain P2947 was exchanged for the P2882-*SS_rtgG_(N6_blpI_)-blpI-HiBiT* PCR product to create strain P3057.

The R6 strains expressing SS_BlpI_(N6_RtgG_)-BlpI-HiBiT were created as follows. A PCR product (P2880-*SS_blpI_(N6_rtgG_)-blpI-HiBiT*) was amplified using primers CW451/CW550 following Gibson assembly of three PCR products amplified using primer pairs CW531/CW595 on R6 template, CW522/CW596 on a gBlocks fragment (IDT) containing *SS_blpI_(N6_rtgG_)-blpI-HiBiT*, and CW597/CW551 on P2967 template. Another PCR product (P2882-*SS_blpI_(N6_rtgG_)-blpI-HiBiT*) was amplified using primers CW451/CW550 following Gibson assembly of three PCR products amplified using primer pairs CW543/CW511 on P2882 template, CW522/CW596 on the *SS_blpI_(N6_rtgG_)*-*blpI-HiBiT* gBlocks fragment from above, and CW597/CW551 on P2967 template. The Sweet Janus+ cassettes in strains P2945, P2949, and P2951 were exchanged for the P2880-*SS_blpI_(N6_rtgG_)-blpI-HiBiT* PCR product to create strains P3081, P3085, and P3087, respectively. The Sweet Janus+ cassette in strain P2947 was exchanged for the P2882-*SS_blpI_(N6_rtgG_)-blpI-HiBiT* PCR product to create strain P3083.

The R6 strains expressing SS_RtgG_[ΔN(2-6)]-BlpI-HiBiT or SS_BlpI-ΔN(2-6)_-BlpI-HiBiT were created as follows. A PCR product (P2882-*SS_rtgG_*[*ΔN(2-6)*]*-blpI-HiBiT*) was amplified using primers CW451/CW550 following Gibson assembly of three PCR products amplified using primer pairs CW543/CW595 on P2882 template, CW522/CW596 on a gBlocks fragment (IDT) containing *SS_rtgG_*[*ΔN(2-6)*]*-blpI-HiBiT*, and CW597/CW551 on P2967 template. Another PCR product (P2882-*SS_blpI_*[*ΔN(2-6)*]*-blpI-HiBiT*) was amplified using primers CW451/CW550 following Gibson assembly of three PCR products amplified using primer pairs CW543/CW511 on P2882 template, CW522/CW596 on a gBlocks fragment (IDT) containing *SS_blpI_*[*ΔN(2-6)*]*-blpI-HiBiT*, and CW597/CW551 on P2967 template. The Sweet Janus+ cassette in strain P2947 was exchanged for the P2882-*SS_rtgG_*[*ΔN(2-6)*]*-blpI-HiBiT* and P2882-*SS_blpI_*[*ΔN(2-6)*]*-blpI-HiBiT* PCR products to create strains P3166 and P3170, respectively.

The R6 strains expressing SS_RtgG_-BlpI-HiBiT with single-residue substitutions in the signal sequence were created as follows. PCR products were amplified using primers CW451/CW550, each following Gibson assembly of two PCR products amplified using the following primer pairs on P2983 template: CW543/CW655 and CW656/CW551 [E(−22)A], CW543/CW654 and CW656/CW551 [L(−21)A], CW543/CW653 and CW656/CW551 [I(−20)A], CW543/CW652 and CW656/CW551 [L(−19)A], and CW543/CW649 and CW651/CW551 [P(−18)M]. The Sweet Janus+ cassette in strain P2947 was exchanged for the above PCR products to create strains P3133, P3135, P3137, P3138, and P3160, respectively.

### Sequencing of the Sp9-BS68 rtg locus

To bridge gaps in the *rtg* locus in the published genome sequence of Sp9-BS68 (11), we amplified the region between *rtgG* and *rtgD2* using primers CW545/CW539 and performed Sanger sequencing using primers CW640, CW665, CW518, CW666, CW668, CW568, and CW593. This allowed us to manually and unambiguously join contigs 306 (ABAB01000022), 342 (ABAB01000053), 99 (ABAB01000059), and 173 (ABAB01000005).

### P_rtgA_-luc lytic luciferase reporter assay

Cells expressing P*_rtgA_*-*luc* reporters were grown in THY, CDM+, or RPMI (Thermo Fisher Scientific, 11875093) ± 1% fetal bovine serum (Thermo Fisher Scientific, 10437028). At OD_620_ 0.2, cells were harvested, pelleted by centrifugation at 6000×*g*, 5 min, 4°C, washed with PBS, pelleted again and resuspended in lysis buffer (50 mM Tris-HCl pH 7.5, 10 mM MgCl_2_, 1 mM DTT, 0.2% Triton X-100). After incubation with lysis buffer for 15 min at room temperature, the lysed samples were mixed with firefly luciferase reaction buffer [50 mM Tris-HCl pH 7.5, 10 mM MgCl_2_, 1 mM DTT, 2 mM firefly luciferin (Thermo Fisher Scientific, 88294), 2 mM ATP, 1 mM coenzyme A] at a 1:1 ratio and luminescence was immediately read with a Synergy HTX plate reader (Biotek) in a white 96-well plate (Costar, 3917). Differences between groups were assessed by ANOVA followed by Tukey’s HSD test, using the aov() and TukeyHSD() functions in R 3.5.1.

### Calculations for rtg, com, and blp activation events

Adapted from ref (4). Analysis was restricted to timepoints at which OD_620_ was greater than or equal to 0.01. The low values of both luminescence and OD_620_ at timepoints before this cell density threshold resulted in low signal-to-noise ratios that made automated analysis difficult. An activation event was defined as the first timepoint at which the activation level remained above a threshold *TT* for at least three consecutive readings. *TT* was defined as a function of cell density as follows:

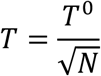

*TT*^0^, threshold constant; *NN*, cell density (OD_620_). This function was chosen because it was the simplest form that empirically yielded a threshold curve with high sensitivity and specificity. The constant *TT*^0^ was empirically determined by manual inspection of activation level curves from pheromone-treated samples as positive controls and curves from non-treated pheromone deletion strains as negative controls. Wells that were already activated at the beginning of the analysis period were left censored and wells that did not activate before they reached their maximum observed cell densities were right censored.

### Peptide secretion assays

Samples for extracellular peptide quantification were read with a Synergy HTX plate reader (Biotek) in a white 96-well plate (Costar, 3917) following addition of HiBiT Extracellular Detection Reagent (Promega, N2421) at a 1:1 ratio. For endpoint assays, each sample was quantified with three technical replicates and read 5 min following reagent addition. For time-course assays, each sample was quantified with one technical replicate and read 1 min following reagent addition. Samples for intracellular peptide quantification were pelleted at 6000×*g*, 5 min, 4°C and resuspended in proteinase K buffer [20 mM MES pH 6.5, 20 mM MgCl_2_, 0.5 M sucrose, 100 µg/mL proteinase K (Fisher Scientific, BP1700)]. A 15-min incubation at 37°C removed residual extracellular peptide by proteinase K digestion. Afterwards, proteinase K was inactivated by addition of 1 mM phenylmethanesulfonyl fluoride (Calbiochem, 7110). Cells were lysed by addition of 1% Triton X-100 followed by incubation at room temperature for 15 min. The lysed samples were then mixed with HiBiT Extracellular Detection Reagent at a 1:1 ratio in a white 96-well plate and read with a Synergy HTX plate reader. Each sample was quantified with three technical replicates and read 5 min following reagent addition.

### Genomic analysis of rtg

Genomes of the Massachusetts pneumococcal isolate collection (BioProject Accession: PRJEB2632) were filtered based on sequence coverage of the predicted location of *rtg*. Three possible upstream flanking genes were defined: a gene coding for a LysM-domain protein (*SPD_0104*), *argG* (*SPD_0110*), and *argH* (*SPD_0111*). Two possible downstream flanking genes were defined: a putative endoRNAse gene (*SPD_0125*; disrupted in D39) and *pspA* (*SPD_0126*). The D39 versions of these genes were BLASTed (megablast profile) against all genomes in the Massachusetts collection. In the case of *pspA*, only the first 250 nucleotides were used so that the choline-binding repeats found later in the gene did not complicate the analysis. All genomes in which at least one upstream flanking gene was found on the same contig as at least one downstream flanking gene were chosen for further analysis (318 in total); all other genomes were discarded. For each genome in the filtered set of 318, the sequence between the two closest flanking genes was extracted and a distance matrix for these sequences representing *rtg* loci was calculated using BIGSdb (12). The set of unique *rtg* genes found in Sp9-BS68 and D39 was used as the reference gene list. Then, a neighbor-joining tree was constructed from the distance matrix using SplitsTree (13). This tree was used to guide manual clustering of the *rtg* loci into different groups based on gene presence and synteny. New unique *rtg* genes found in the course of this analysis were added to the initial set of genes from Sp9-BS68 and D39 and the updated set was BLASTed (blastn profile) against all 616 genomes in the Massachusetts collection to determine how many strains in the full collection encoded *rtg*. Genomes with a hit (≥ 80% query coverage, ≥ 70% sequence identity) for at least one *rtg* gene were considered to be *rtg*-positive.

### Calculations for mouse colonization assays

The competitive nasal colonization performed for this publication was part of an approved protocol authorized by the University of Michigan IACUC. Ten-fold serial dilutions of PBS suspensions used to inoculate mice were spot plated (5 µL per spot, 3 replicates per sample) on tryptic soy agar (TSA) plates supplemented with 5 µg/mL catalase and containing either neomycin (selects for all pneumococcus), spectinomycin, or gentamicin. Plates were incubated at 37°C, 5% CO_2_ overnight. Afterwards, colony counts were obtained. To obtain colonization density data from singly inoculated mice, nasal washes were quantified in the same manner as described above, and counts from the neomycin plates were used. When attempting to quantify colonization density from mice inoculated with two competing strains, we noticed that the growth of the gentamicin-resistant strain on gentamicin-containing plates were inhibited by an unknown factor from the nasal wash. This occurred during both trials of the colonization assay and occurred in singly colonized mice who received the gentamicin resistant strain. This inhibition was not seen for the gentamicin-resistant strain growing on neomycin-containing plates from the singly inoculated mice. No inhibition was seen for either of the spectinomycin-resistant strains growing on either spectinomycin- or neomycin-containing plates. This inhibition was also not seen for any strain growing on any plate during plating of the inoculants. This phenomenon prevented us from obtaining accurate counts of the gentamicin-resistant strain from co-inoculated mice using the spot-plating method. However, since the inhibition was restricted to only gentamicin-containing plates, we developed an alternative method for quantifying the ratio of spectinomycin-resistant to gentamicin-resistant colonies isolated from co-inoculated mice using the samples plated on neomycin-containing plates. For each mouse, we picked 40-100 colonies from the spots on the neomycin-containing plates, grew them in THY + 5 µg/mL catalase at 37°C for 6-8 hours, then replicate-plated the cultures on TSA plates supplemented with 5 µg/mL catalase and containing either spectinomycin or gentamicin. After an overnight incubation at 37°C with 5% CO_2_, each culture was scored for growth on spectinomycin or gentamicin. Cultures that showed growth on both antibiotics were excluded from further analysis. If none of the cultures from a mouse grew on a particular antibiotic, these data points were treated as being below the limit of detection and a count of 1 was used instead of 0. For each mouse, the output ratio was defined to be the count of cultures showing spectinomycin-resistant growth divided by the count of cultures showing gentamicin-resistant growth. We confirmed that neither of the spectinomycin-resistant strains could inhibit growth of the gentamicin-resistant strain or vice versa during growth on neomycin-containing plates or in co-culture in THY by repeating the above method on spot-plated samples from mixed cultures prepared in an identical fashion to those used to inoculate mice. Ratios obtained from the replicate-plating method agreed closely with ratios obtained from traditional counting of colonies on spectinomycin- and gentamicin-containing plates. The final competitive index calculation was performed using the following formula:

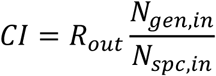

*CI*, competitive index; *R_out_*, output ratio; *N_gen,in_*, input CFU density of gentamicin-resistant strain; *N_spc,in_*, input CFU density of spectinomycin-resistant strain.

---

**Figure S1.**
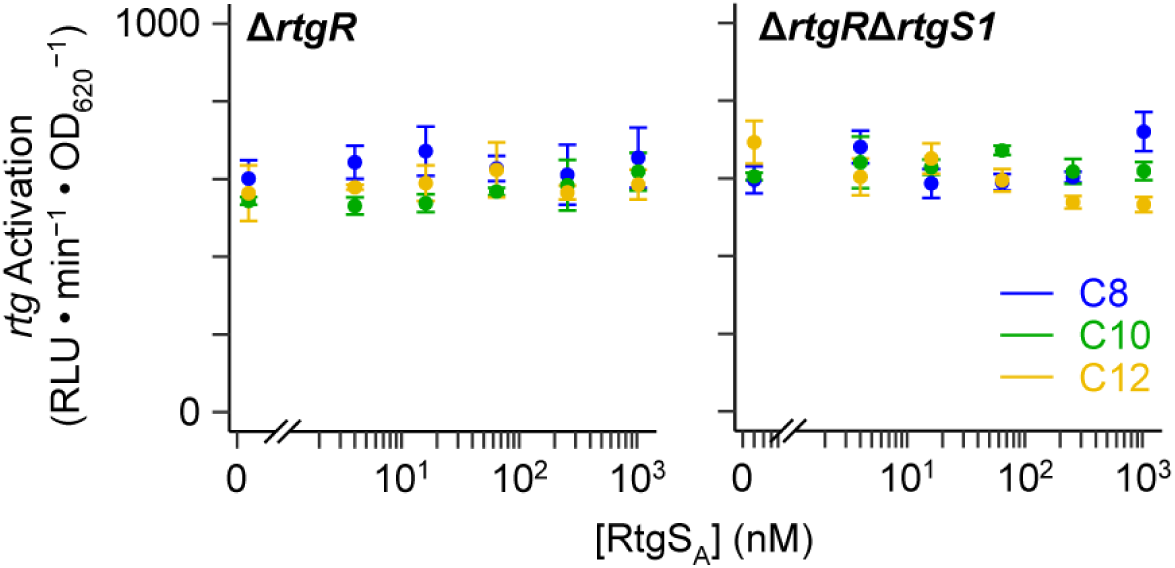
RtgS-induced *rtg* activation requires RtgR. Sp9-BS68 Δ*rtgR* and Δ*rtgR*Δ*rtgS1* P*_rtgS1_*-*luc* reporters were grown in CDM+ to OD_620_ 0.02 and treated with synthetic RtgS_A_ fragments. Response was defined as the maximum observed P*_rtgS1_* activity within 60 min of treatment. Plotted data points represent mean ± S.E. of 3 independent experiments.

**Figure S2.**
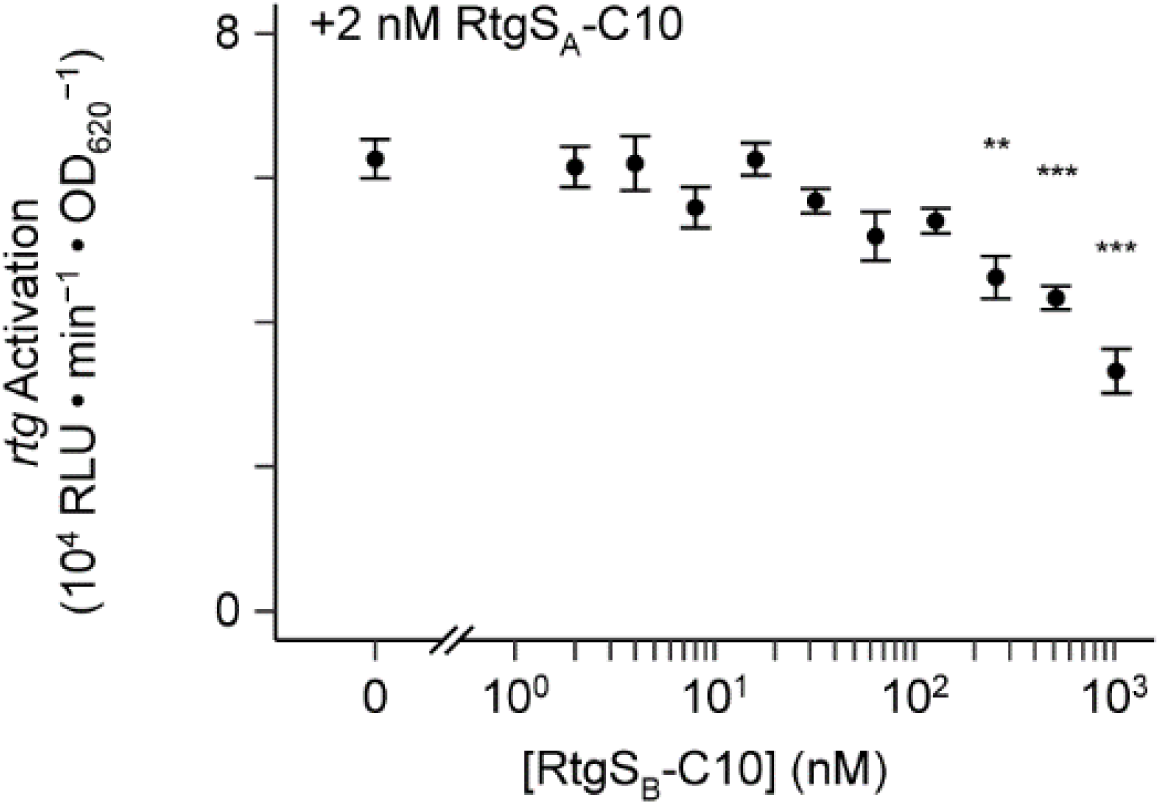
High concentrations of RtgS_B_-C10 antagonize RtgS_A_-C10. D39 Δ*rtgS1*Δ*rtgS2* P*_rtgS1_*-*luc* reporters were grown in CDM+ to OD_620_ 0.02 and treated simultaneously with 2 nM RtgS_A_-C10 and various concentrations of RtgS_B_-C10. Response was defined as the maximum observed P*_rtgS1_* activity within 120 min of treatment. Plotted data points represent mean ± S.E. of 3 independent experiments. Statistics: comparisons vs. 0 nM RtgS_B_-C10; ** p < 0.01, *** p < 0.001; ANOVA with Dunnett’s correction for multiple comparisons.

**Figure S3.**
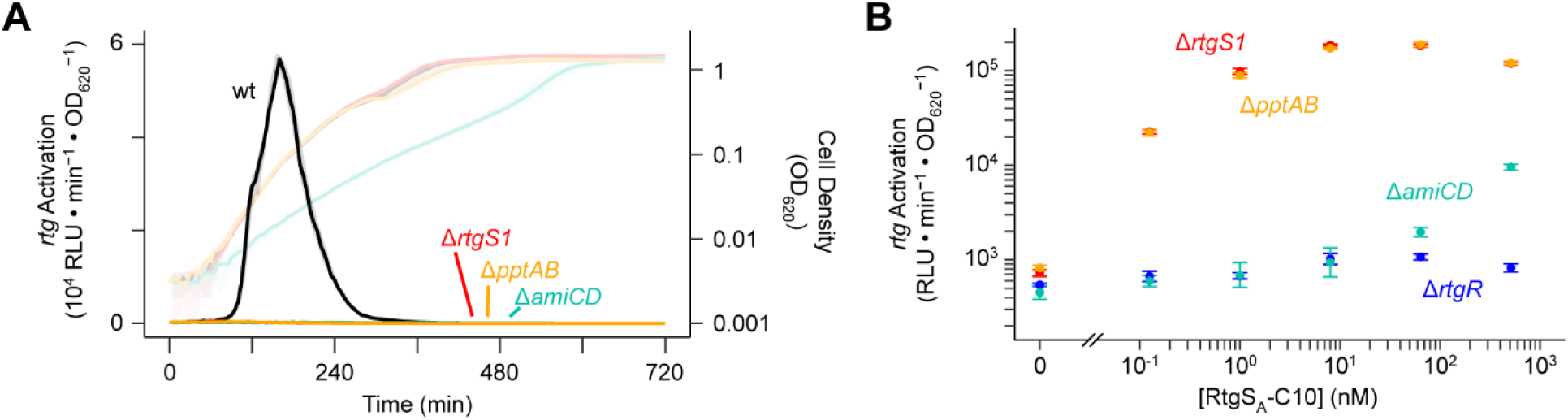
Ami and PptAB are required for RtgR/S signaling. **(A)** Both Ami and PptAB are required for natural *rtg* autoinduction. Sp9-BS68 P*_rtgS1_*-*luc* reporters were grown in CDM+ and monitored for *rtg* activation (dark, left y axis) and cell density (light, right y axis). Plots show median (line) and 25% to 75% quantiles (shading) of 30 wells per strain pooled from 3 independent experiments. **(B)** Exogenous RtgS treatment rescues *rtg* activation defect in the Δ*pptAB* mutant but not the Δ*amiCD* mutant. Sp9-BS68 P*_rtgS1_*-*luc* reporters were grown in CDM+ to OD_620_ 0.02 and treated with RtgS_A_-C10. Response was defined as the maximum observed P*_rtgS1_* activity within 60 min of treatment. Plotted data points represent mean ± S.E. of 3 independent experiments.

**Figure S4.**
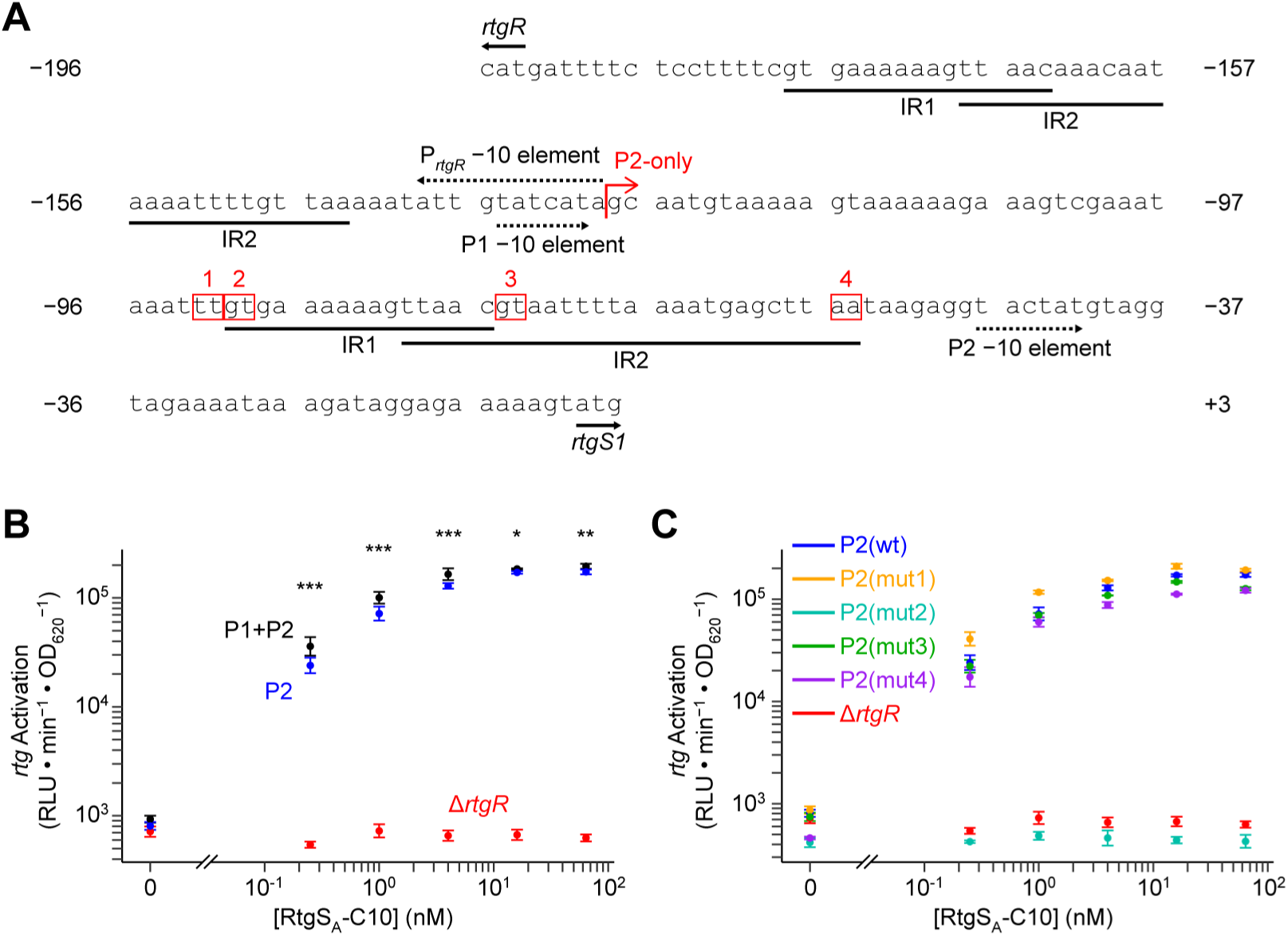
Two promoters, P1 and P2, both with putative RtgR binding sites, contribute to *rtgS1* expression. **(A)** Promoter region of *rtgR* and *rtgS1* in Sp9-BS68. Inverted repeats (IR) are underlined. −10 elements are marked with dotted arrows. Start sites of *rtgR* and *rtgS1* are marked with solid arrows. Red bent arrow indicates span of P2 promoter construct used in panel B. Numbered, red boxes indicate mutated nucleotides used in panel C. **(B)** P1 and P2 each partially contribute to *rtgS1* expression. Sp9-BS68 Δ*rtgS1* P*_rtgS1_*-*luc* reporters were grown in CDM+ to OD_620_ 0.02 and treated with synthetic RtgS_A_-C10. Response was defined as the maximum observed P*_rtgS1_* activity within 60 min of treatment. Plotted data points represent mean ± S.E. of 3 independent experiments. Statistics: comparisons between P1+P2 and P2 reporters at each dose; * p < 0.05, ** p < 0.01, *** p < 0.001; ANOVA followed by Holm-corrected post-tests. **(C)** IR1 is required for RtgS-induced *rtg* activation. Sp9-BS68 Δ*rtgS1* P*_rtgS1_*-*luc* reporters were grown in CDM+ to OD_620_ 0.02 and treated with synthetic RtgS_A_-C10. Except for the Δ*rtgR* strain, *luc* in each reporter was fused to P2 only. Response was defined as the maximum observed P*_rtgS1_* activity within 60 min of treatment. Plotted data points represent mean ± S.E. of 3 independent experiments.

**Figure S5.**
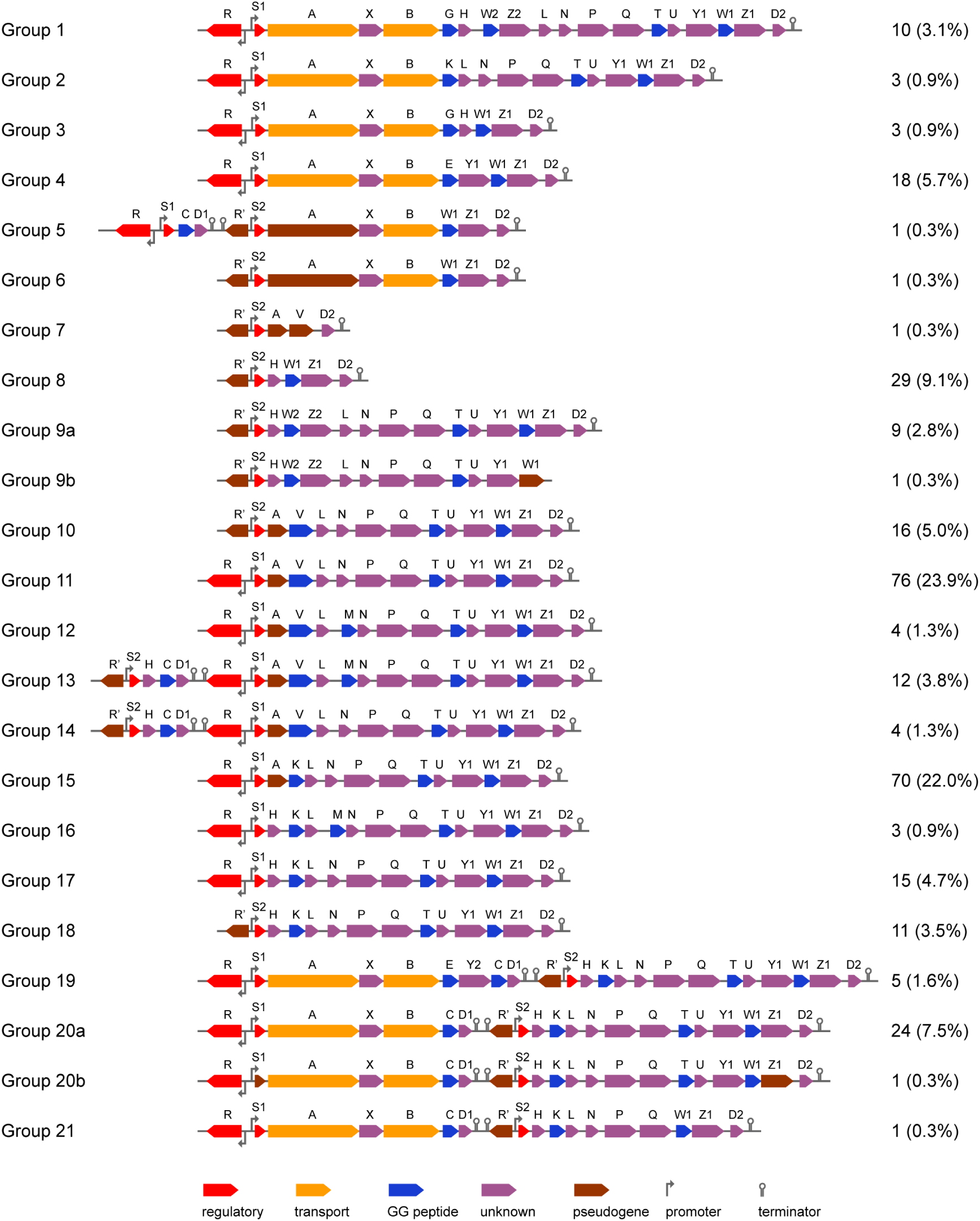
Diversity of fully sequenced *rtg* loci in the Massachusetts pneumococcal isolate collection. *rtg* loci from a filtered set of 318 strains from the Massachusetts isolate collection with gapless sequence coverage of the *rtg* genomic locale were clustered into 23 groups based on gene content and synteny. The organization of *rtg* in each of these groups is depicted here. Block arrows represent genes and are color-coded according to their predicted function/classification. Genes not explicitly marked as pseudogenes may still be present as pseudogenes in a subset of strains within a group. The number and percentage of strains (out of 318) belonging to each group is displayed on the right.

**Figure S6.**
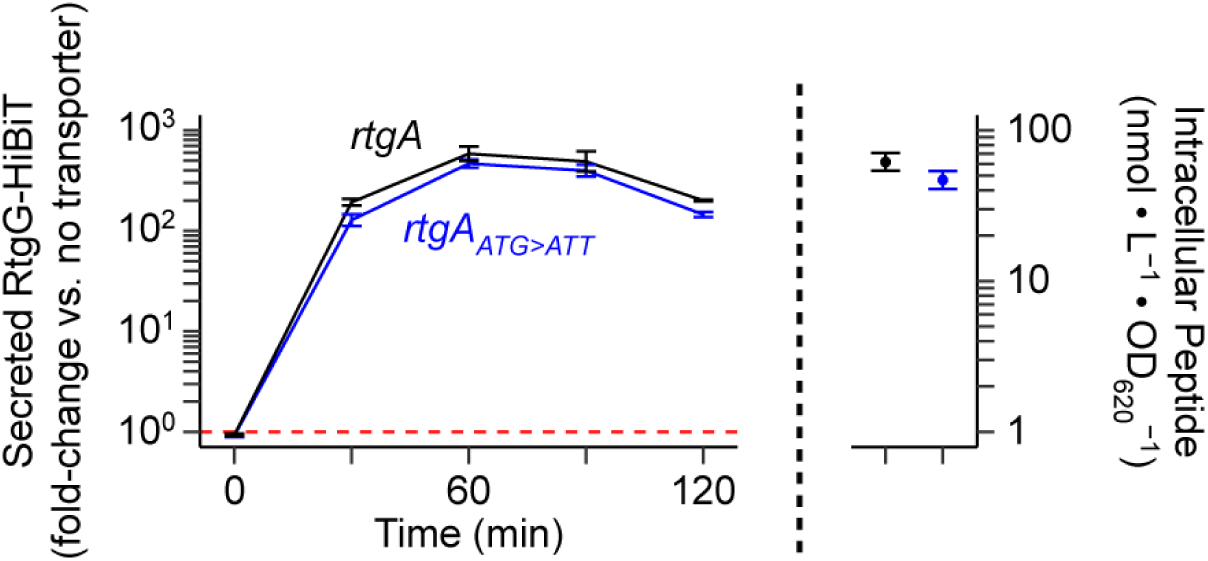
Strains with the *rtgA_ATG>ATT_* mutation still produce functional RtgAB. R6 ComAB^−^/BlpAB^−^/RtgAB^−^ and RtgAB^+^ strains with (blue) or without (black) the ATG>ATT mutation in *rtgA* expressing RtgG-HiBiT were grown in CDM+ to OD_620_ 0.05 and treated with 200 ng/mL CSP, 200 ng/mL BlpC, and 20 nM RtgS_A_-C10. Samples were taken every 30 min and extracellular HiBiT signal was quantified (left). Data are presented as fold-change values relative to the ComAB^−^/BlpAB^−^/RtgAB^−^ control. Red, dashed line represents fold-change = 1 (no difference vs. the control). At the 120-min timepoint, intracellular peptide content was also quantified (right). Plots show mean ± S.E. of 3 independent experiments.

**Figure S7.**
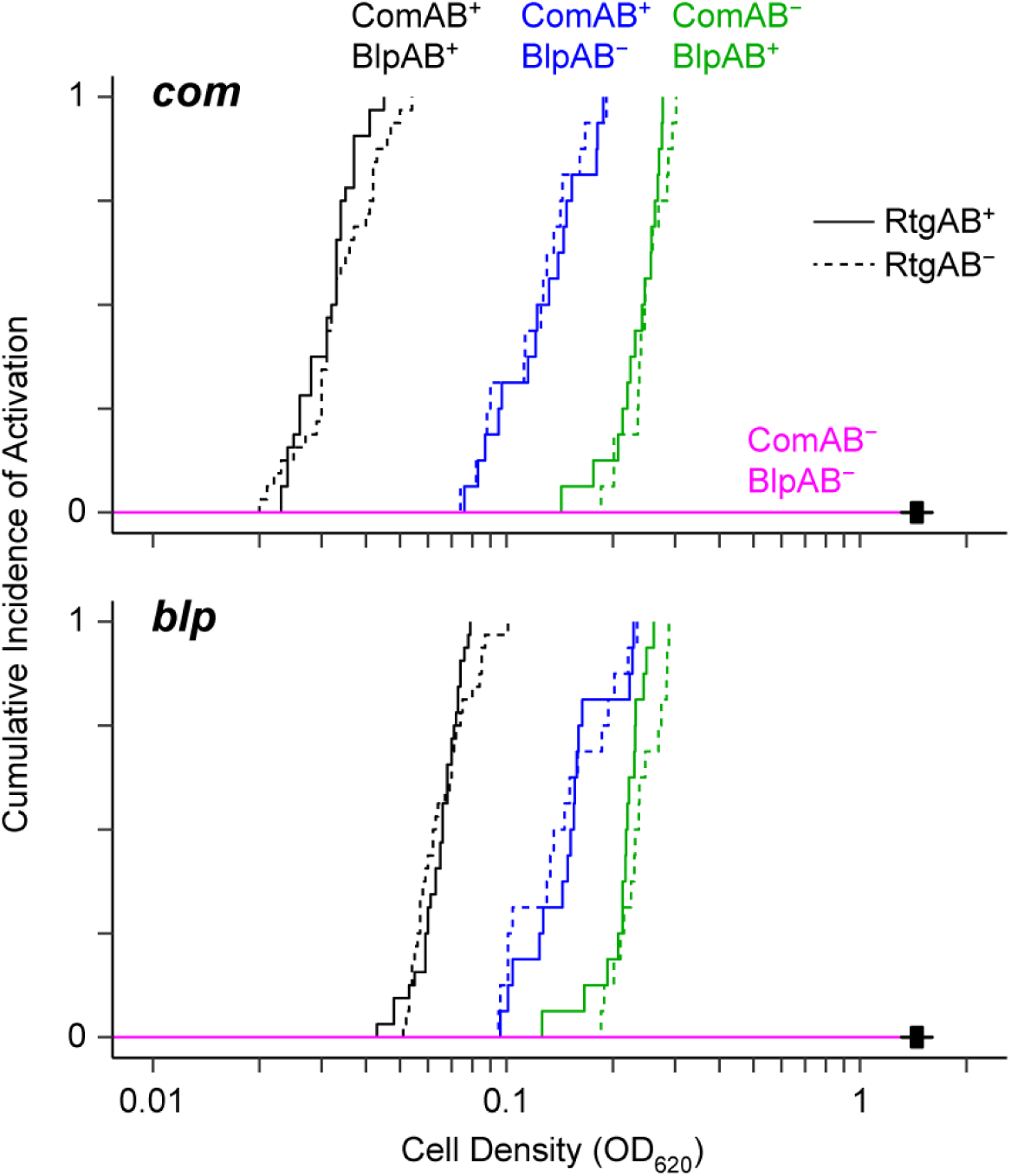
RtgAB does not affect the timing of *com* or *blp* activation during growth in CDM+. D39 P*_comA_*-*Nluc* P_BIR_-*RFluc* dual reporters were grown in CDM+ and monitored for *com* (top) and *blp* (bottom) activation. Data were fit to the Kaplan-Meier estimator. Wells that did not experience an activation event before cells reached their maximum density were censored (crosses). N = 32 (ComAB^+^/BlpAB^+^ strains) or 16 (all other strains) wells per strain pooled from 4 independent experiments. Statistics: all comparisons between RtgAB^+^ and RtgAB^−^ strains with identical ComAB/BlpAB phenotypes (except in the ComAB^−^/BlpAB^−^ background, for which all data points were censored) were not significant (p ≥ 0.05); log-rank test with Holm correction.

**Figure S8.**
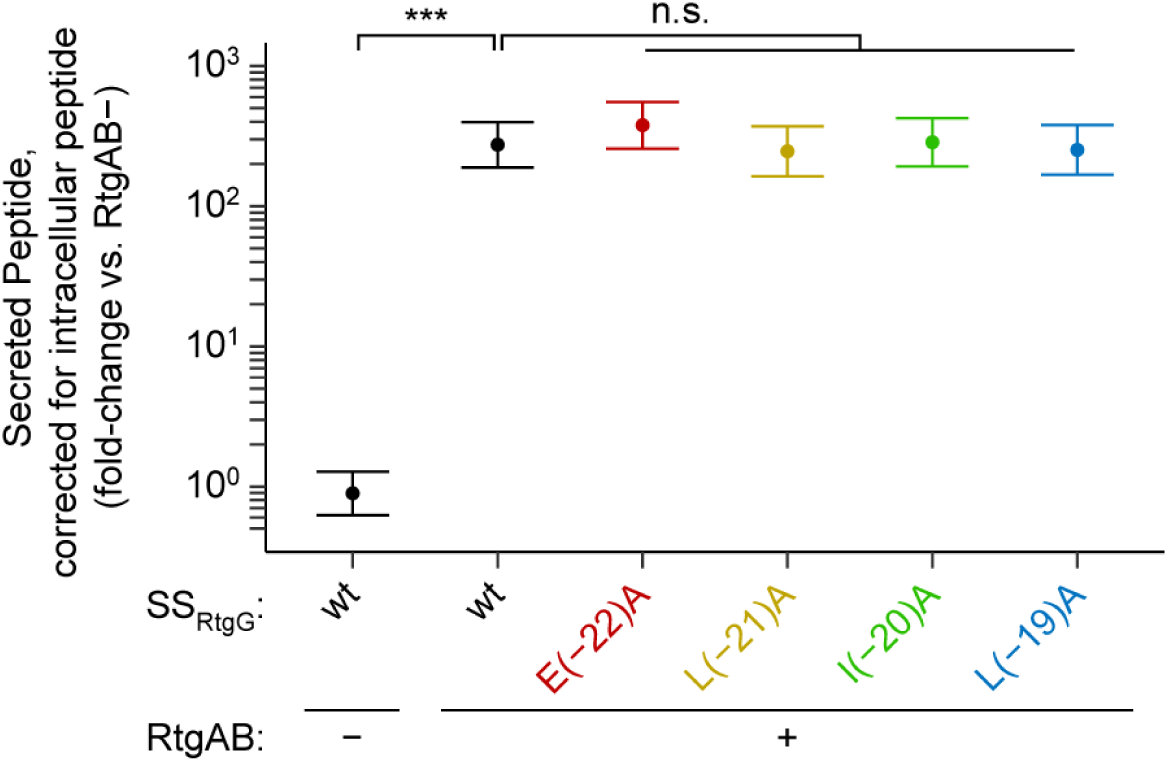
Single alanine substitutions in the N-terminal region of SS_RtgG_ do not decrease secretion through RtgAB. R6 ComAB^−^/BlpAB^−^/RtgAB^−^ and RtgAB^+^ strains expressing mutated SS_RtgG_ fused to BlpI-HiBiT cargo peptide were grown in CDM+ to OD_620_ 0.05 and treated with 200 ng/mL CSP, 200 ng/mL BlpC, and 20 nM RtgS_A_-C10. Extracellular and intracellular HiBiT signals were quantified at 60 min post-treatment. Corrected secreted peptide levels were calculated by normalizing extracellular signals to intracellular signals. Data are presented as fold-change values relative to the ComAB^−^/BlpAB^−^/RtgAB^−^ control strain with wild-type SS_RtgG_. Plots show mean ± S.E. of 3 independent experiments. Statistics: n.s. not significant, *** p < 0.001; ANOVA with Dunnett’s post-test.

**Figure S9.**
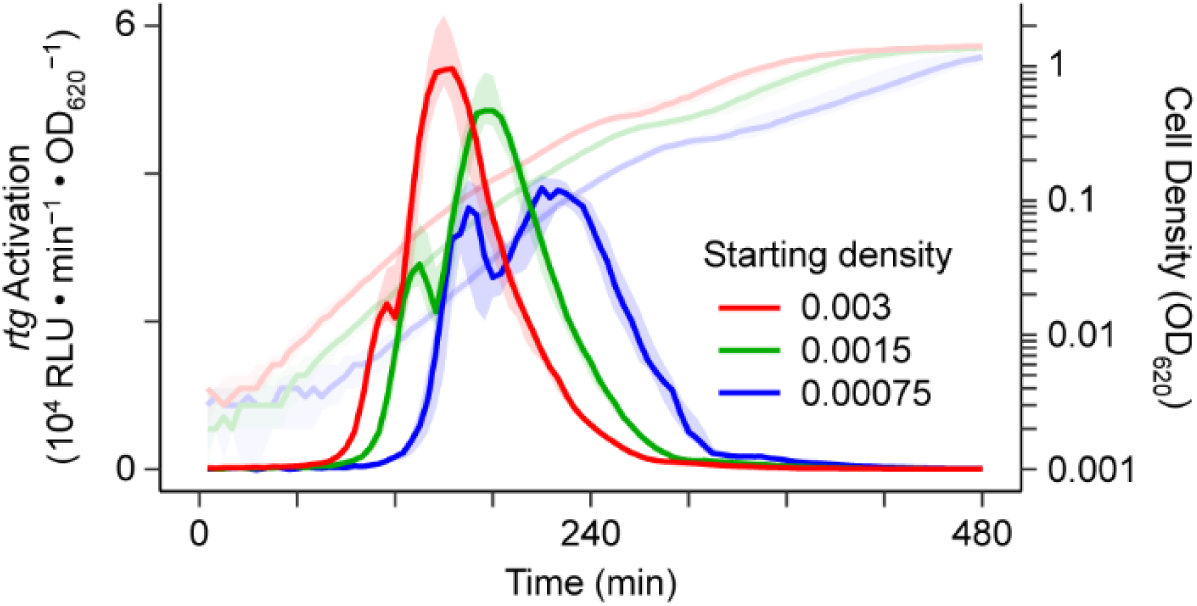
Timing of *rtg* activation depends on cell density. The wildtype Sp9-BS68 P*_rtgS1_*-*luc* reporter was inoculated into CDM+ at different starting densities, grown, and monitored for *rtg* activation (dark, left y axis) and cell density (light, right y axis). Plots show median (line) and 25% to 75% quantiles (shading). N = 30 wells pooled from 3 independent experiments for each of the starting densities of 0.003 and 0.0015. N = 29 wells pooled from 3 independent experiments for starting density of 0.00075; data from one well was discarded due to lack of growth.

**Table S1.** Rgg regulators associated with WxW-motif pheromones.

| Species | Strain | Rgg Gene | % Identity (% Similarity)<br>with RtgR<sub>Sp9-BS68</sub> | Predicted Pheromone |
| --- | --- | --- | --- | --- |
| *Streptococcus pneumoniae* | D39 | SPD_1518 | 42.7% (62.2%) | MGFKKYLKNLPKNSGFLIWSWIQLIWFETWFWG |
| *Streptococcus sanguinis* | SK36 | SSA_2251 | 45.5% (70.5%) | MKKIVYNLILLAVTSIVTTSVFPWWLWW |

**Table S2.**
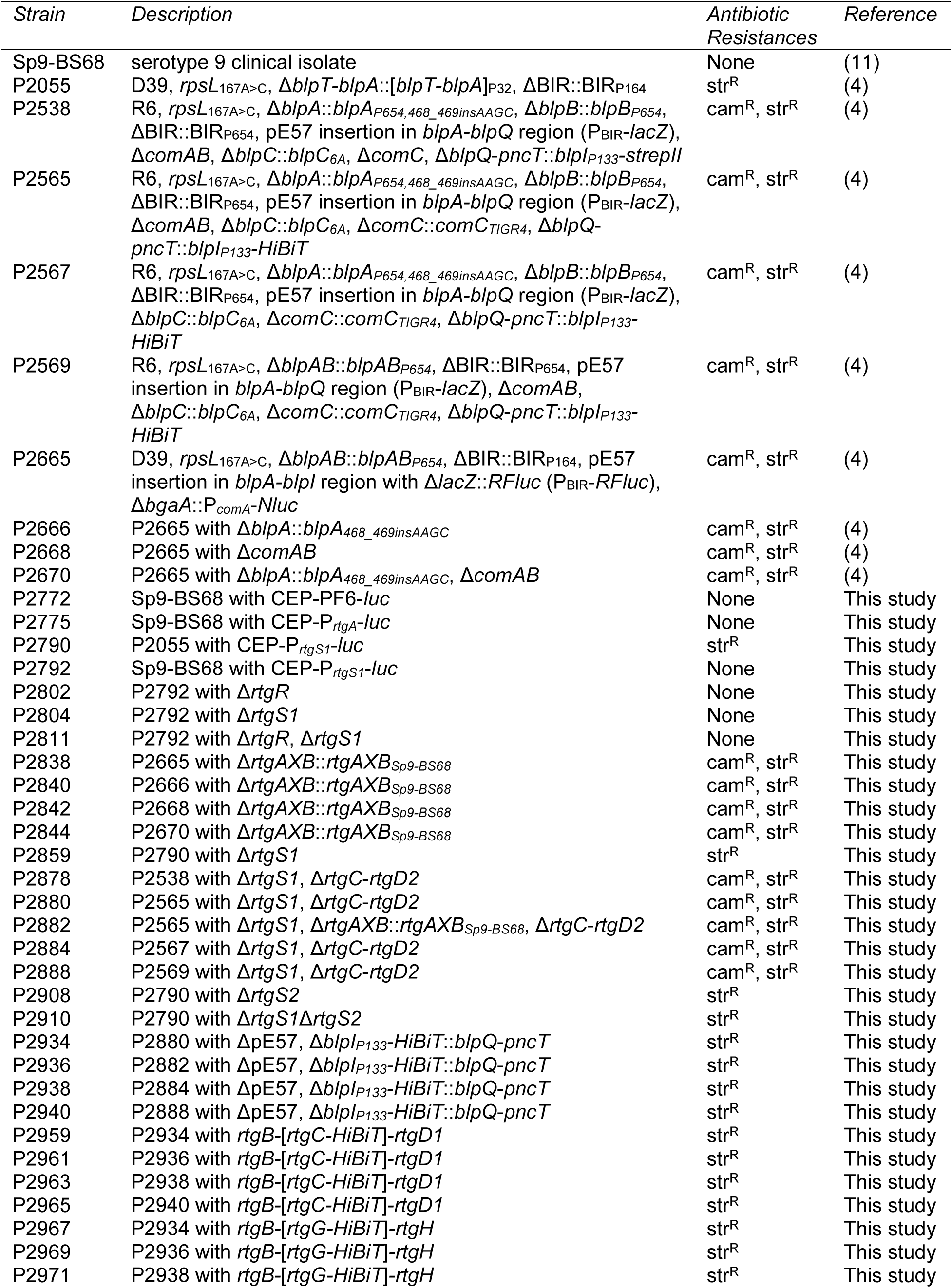
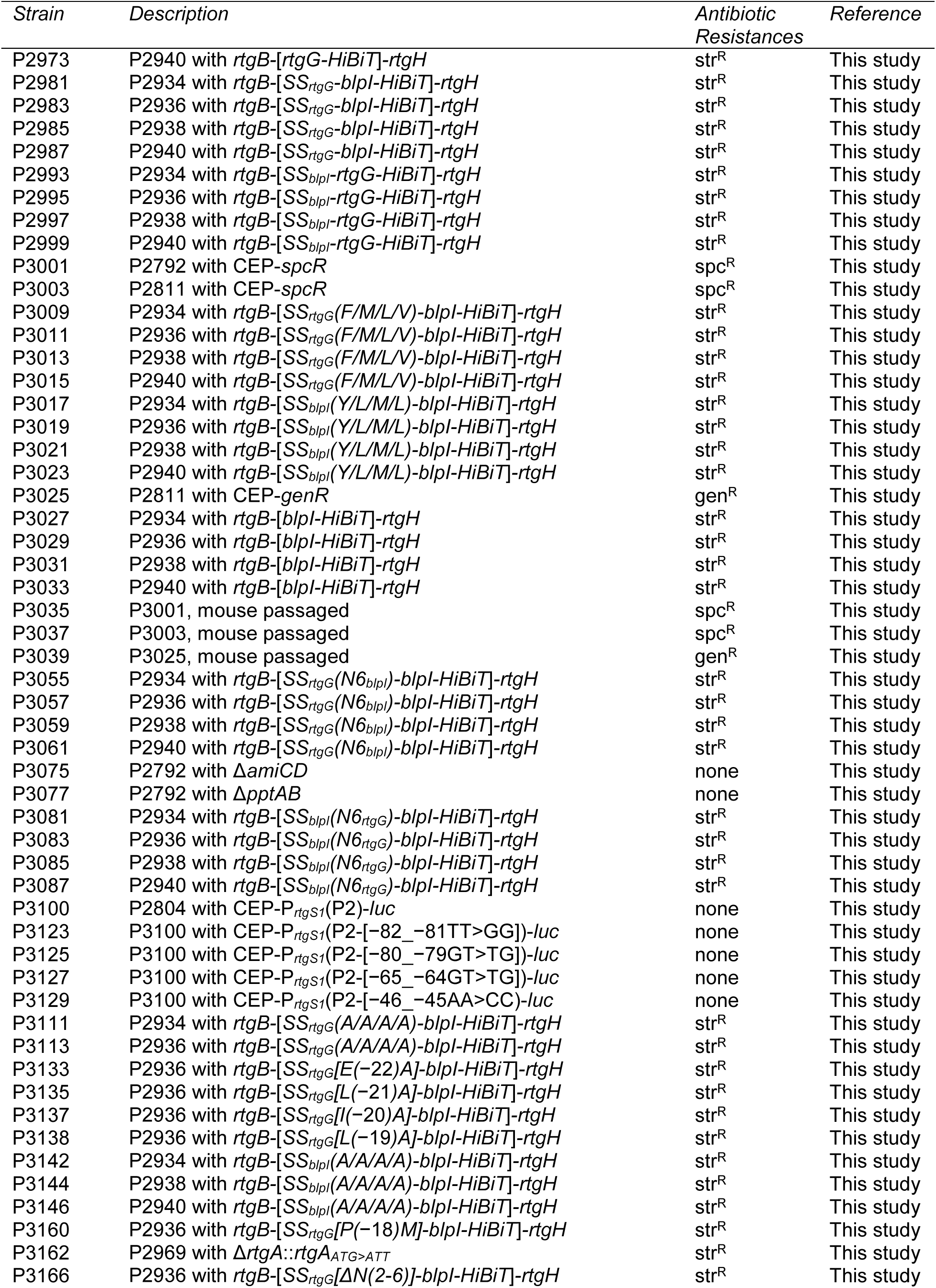
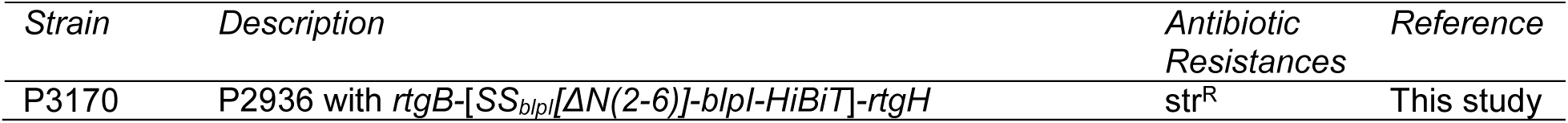
Strain list.

**Table S3.**
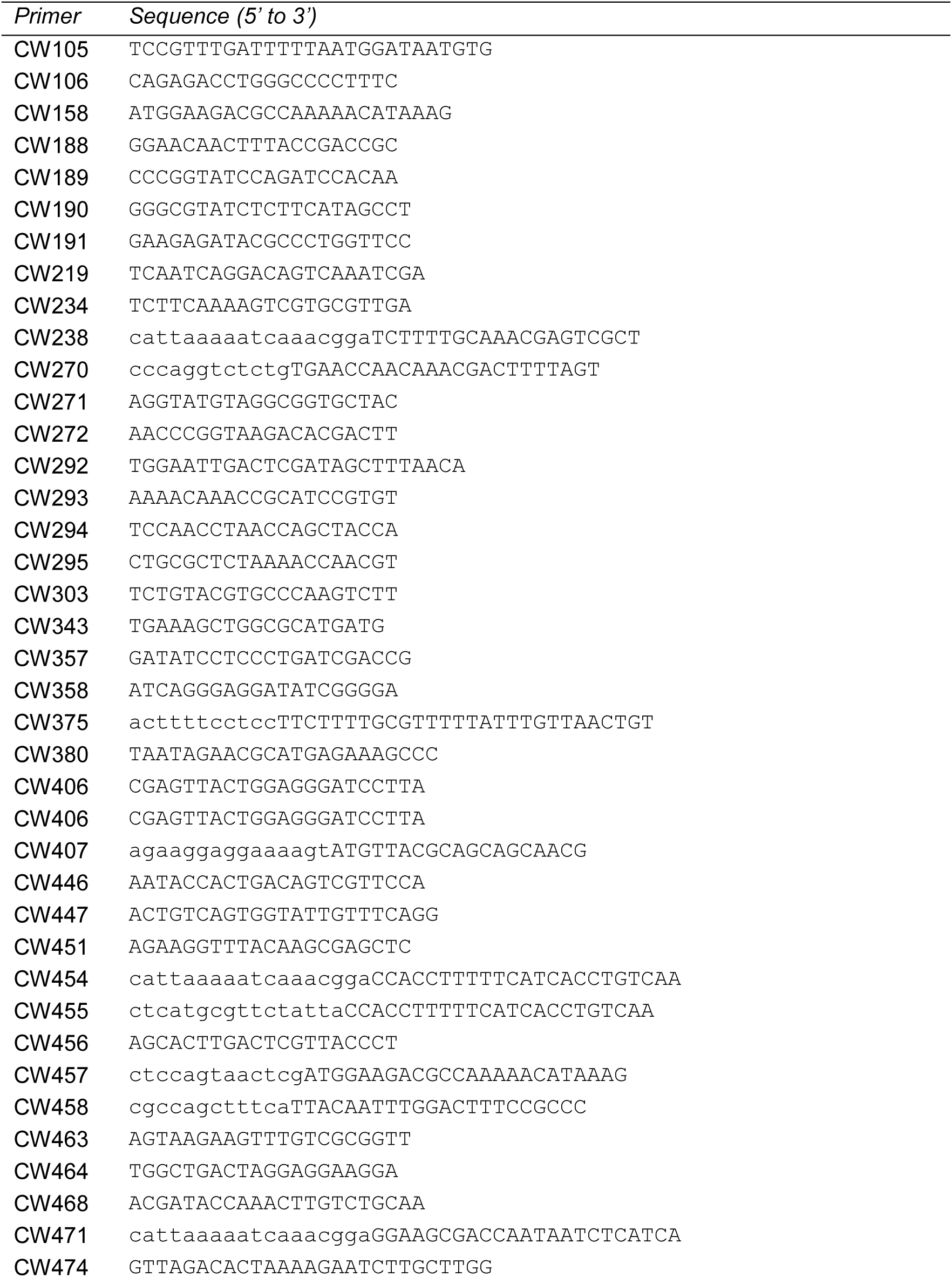
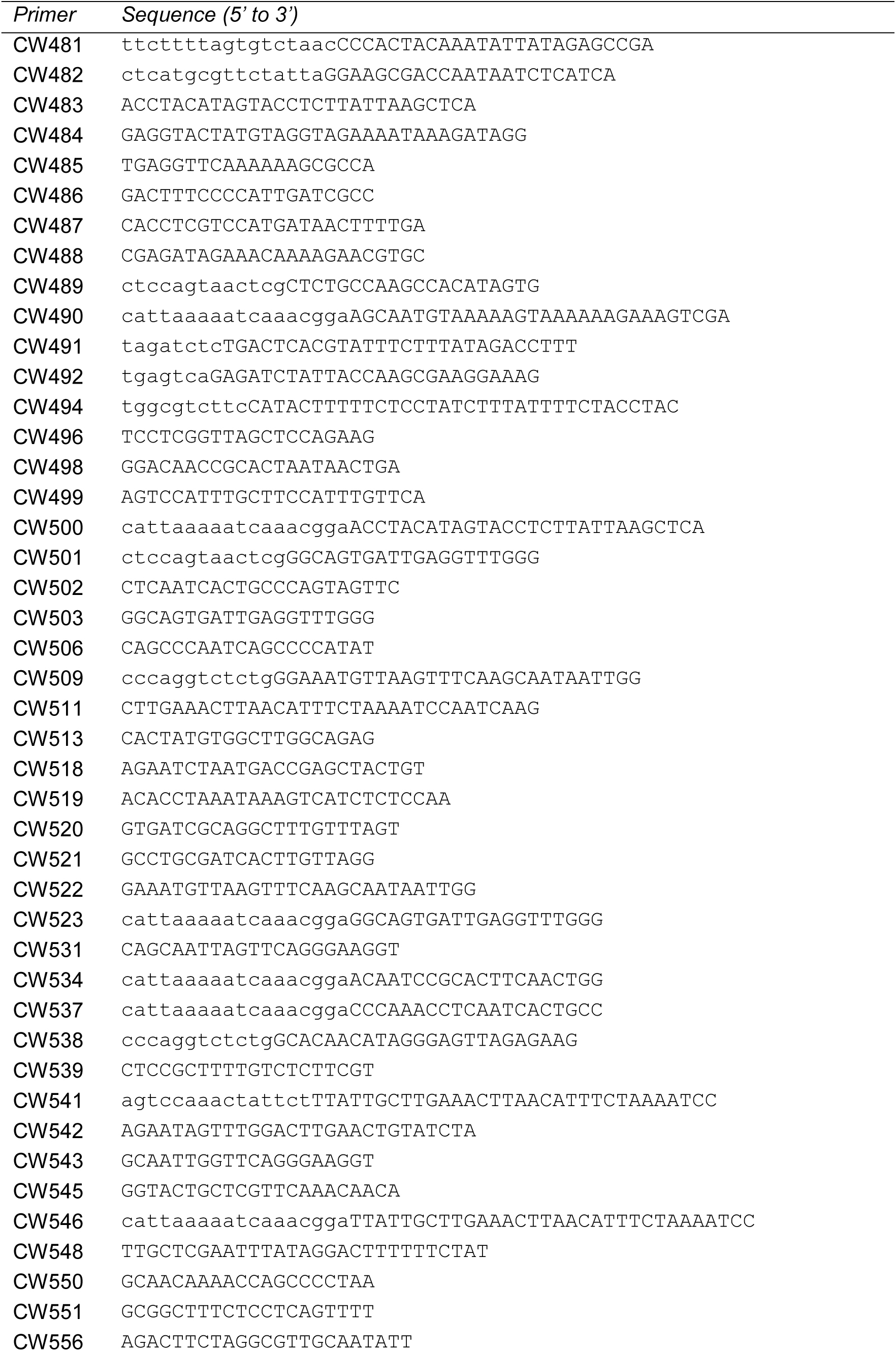
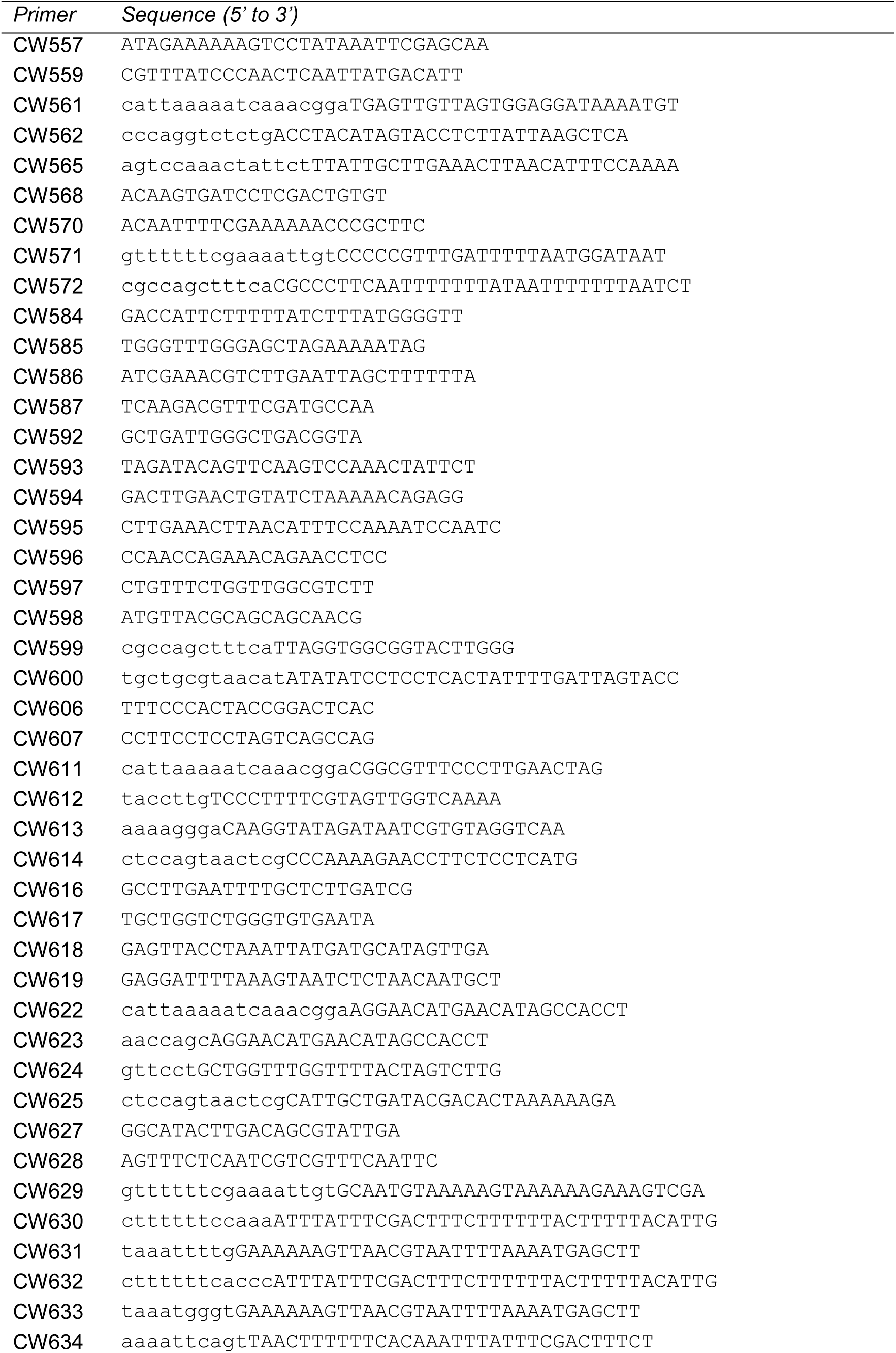
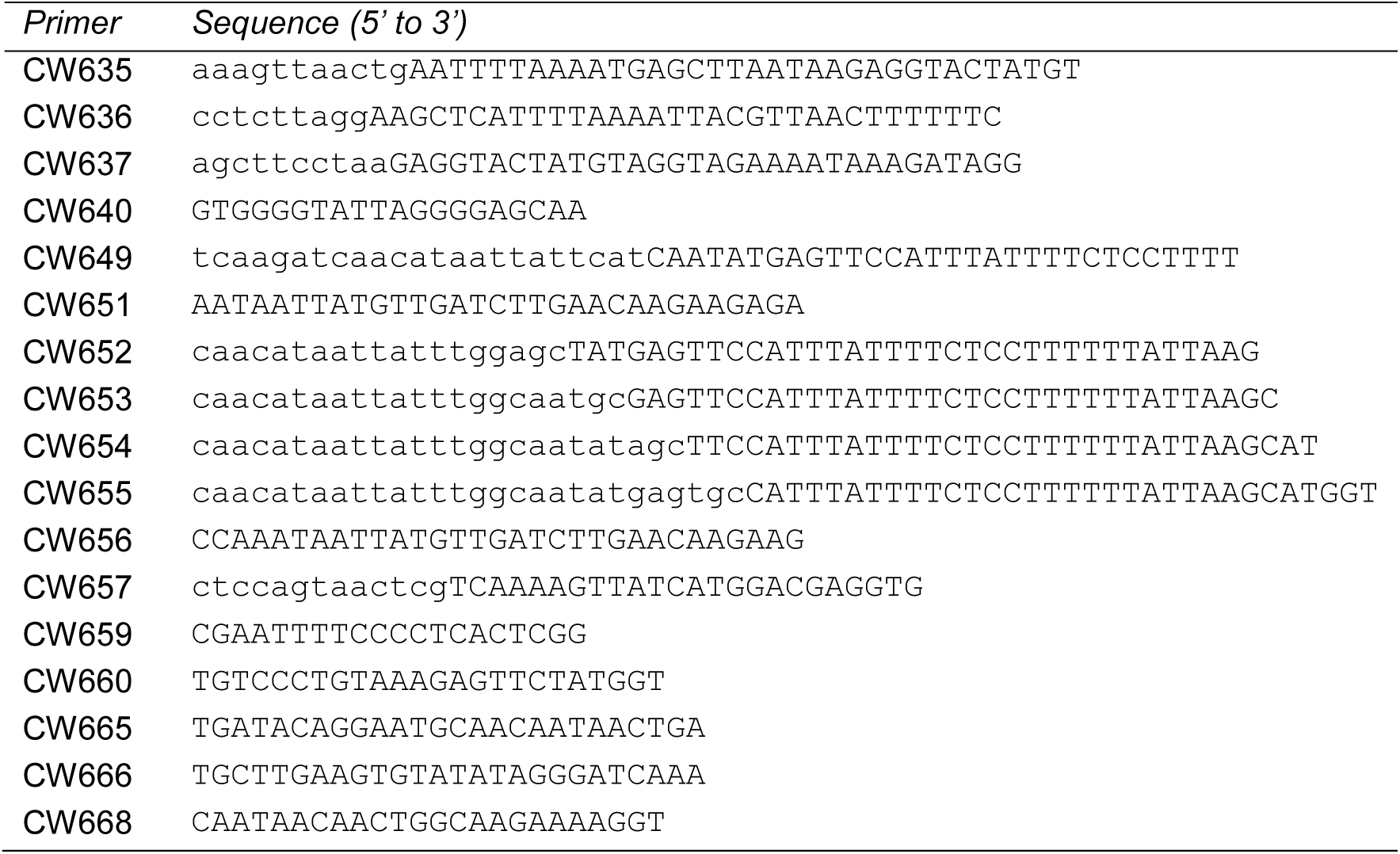
Primer list.

